# Species turnover dominates plant trait variation along continental climate gradients

**DOI:** 10.64898/2026.11.22.767446

**Authors:** Travis G. Britton, Isaac R. Towers, Daniel S. Falster, Laura J. Williams, Rachael V. Gallagher, Ian J. Wright

## Abstract

Variation in plant functional traits along climate gradients chiefly arises from two processes: turnover of species with different trait values, and trait adjustment within species across environments. Yet, the relative contributions of these processes remain poorly quantified, limiting our ability to predict how traits, species distributions and, in turn, ecosystem properties will respond to climate change. Across seven functional traits and 1867 plant taxa from continental Australia, we provide a unified assessment of among-versus within-species contributions across multiple dimensions of plant function. Species turnover emerged as the dominant process driving trait variation along both precipitation and temperature gradients.

Nevertheless, the contribution of within-species adjustment to climate differed substantially among traits and was generally greater in non-woody than woody taxa, these patterns being most clearly expressed along precipitation gradients. Our findings imply that plant traits will track changing climatic conditions primarily through species replacement, particularly for traits such as wood density and seed mass. Species may also respond *in situ* through adjustments in leaf physiology and morphology and, in woody taxa, through shifts in relative allocation between leaves and stem tissues. This knowledge has potential to improve predictions of the structure and function of future plant communities.

## MAIN TEXT

Many plant functional traits exhibit predictable patterns across climate gradients (Givnish 1987, Smith et al. 2019, Towers et al. 2023). These patterns are typically reported in large-scale trait–climate analyses based on species or species-by-site trait means, implicitly integrating variation arising from species turnover (interspecific trait variation) and from within-species trait adjustments via plasticity or genetic differentiation along climate gradients (intraspecific variation; Cornwell and Ackerly 2009, Lepš et al. 2011). Which source of variation dominates in any given situation has different implications for vegetation responses to environmental change (Anderegg 2015). Where trait variation along climate gradients is driven primarily by within-species sources, this suggests a greater potential for rapid *in situ* response via plastic changes within existing species (Nicotra et al. 2010). By contrast, trait–climate relationships dominated by species turnover effects are expected to change more slowly, constrained by the pace at which replacement species disperse and establish under new conditions (Ackerly 2003). Therefore, understanding the relative within- and among-species contributions to trait–climate relationships across trait types is required to robustly predict vegetation responses to climate change.

Few studies have partitioned trait–climate relationships into among- and within-species components, and most have considered a single trait, tissue type or biome (Cornwell and Ackerly 2009, Dwyer et al. 2014, Bjorkman et al. 2018, Sandel and Low 2019, Dong et al. 2020, Tian et al. 2024). Consequently, we lack a general understanding of how these contributions vary across functional traits and broad climate gradients. Available evidence also suggests that their relative importance varies among traits and ecosystems. Species turnover dominated leaf mass per area (LMA; or its inverse, specific leaf area, SLA) variation along environmental gradients in Western Australian wheatbelt and Australian rangeland communities (Dwyer et al. 2014, Dong et al. 2020), whereas turnover and within-species variation contributed similarly to SLA variation along a precipitation gradient in Californian grasslands (Sandel and Low 2019). Within-species contributions have been reported as greater for either LMA or leaf size (Cornwell and Ackerly 2009, Sandel and Low 2019, Dong et al. 2020), and as ranging from relatively low to moderate or high for leaf physiological and nutrient traits (Bjorkman et al. 2018, Dong et al. 2020, Tian et al. 2024). Substantial within-species contributions to wood density variation have also been demonstrated at a global scale (Fischer et al. 2026), yet comparable global-scale analyses of leaf trait variation are lacking. A consistent analysis spanning multiple functional traits and broad climate gradients is therefore needed to identify general patterns in the relative importance of among- and within-species components in shaping climate-related trait variation.

Species with different life histories may vary in how their traits respond to climate gradients and in the relative importance of among-versus within-species trait variation. A major functional contrast is between woody and non-woody species (Díaz et al. 2016), which can show differing – and sometimes opposing – trait–climate relationships (Šímová et al. 2018, Towers et al. 2023). In general, trait–climate sensitivity tends to be lower in herbaceous plants, presumably because non woody plants typically exhibit strategies to avoid (rather than tolerate) harsh conditions, including occupying favourable microhabitats in the understorey or retreating into the seedbank in unfavourable seasons or years (Dwyer and Erickson 2016, Šímová et al. 2018). However, where trait–climate relationships are present, within-species variation may be stronger in herbaceous taxa owing to their high phenotypic plasticity, fast generation time and lower dispersal distance (Thomson et al. 2011), which can yield greater local adaptation to prevailing conditions. If this were the case, it would suggest that non-woody taxa have a greater propensity to track changing climatic conditions within species. However, it remains unclear whether this expectation is borne out empirically across macro-scale climatic gradients.

Here, we harness an Australia-wide plant trait database (AusTraits; Falster et al. 2021) to quantify the relative contributions of species turnover and within-species trait adjustments to climate-driven variation (Fig. 1) in seven leaf, stem, reproductive and biomass allocation traits along two key climate axes: mean annual precipitation (MAP) and mean annual temperature (MAT). MAP and MAT variation are largely orthogonal on the Australian continent (Fig. S1), allowing their effects on trait variation – and its underlying sources – to be evaluated largely independently. We focus on ecologically important traits that are known to vary predictably along these climate gradients; expected directions of response (and their rationale) are outlined in Table 1 (Schulze et al. 2006, Moles et al. 2014, Wright et al. 2017, Rumman et al. 2018, Rosas et al. 2019, Dong et al. 2020, Anderegg et al. 2021, Kambach et al. 2023, Towers et al. 2023, Wang et al. 2023). Further, we developed trait-specific hypotheses regarding the capacity for within-species adjustment along climate gradients, based on differences in physiological versus morphological traits, integration across organs (leaves, seeds, wood), and tissue longevity (Table 1; conceptual scenarios shown in Fig. 1). Analyses were conducted across all taxa and separately for woody and non-woody (herbaceous) plants. This allowed us to test the hypothesis that within-species contributions would be lower for woody relative to non-woody taxa on the basis that their longer lifespans should result in stronger selection for traits optimised to long-term climate, coupled with possibly lower trait plasticity due to investment in longer-lived tissues.

**Figure 1.**
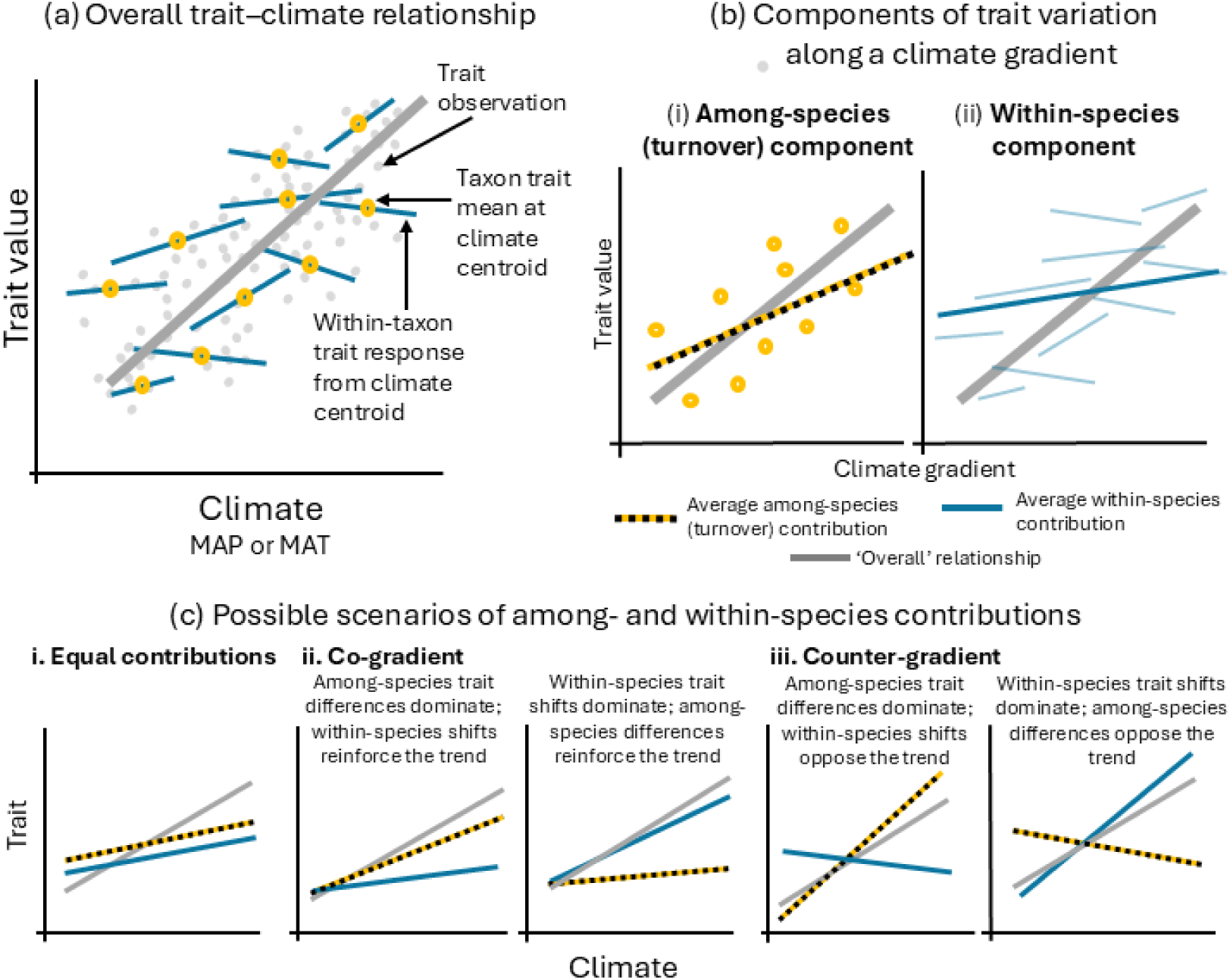
Conceptual framework for partitioning trait–climate relationships into among- and within-species components. (a) The overall trait–climate relationship emerges from the combination of species turnover along the climate gradient (here, MAT or MAP) and within-species trait adjustment. Grey points show individual trait observations; yellow points represent species mean trait values at each taxon’s climate centroid; blue segments indicate within-taxon trait responses relative to deviations from that centroid; the grey line represents the overall trait–climate relationship. (b) The overall relationship can be decomposed into two components: (i) an among-species (turnover) component (the yellow and black dashed line), reflecting the change in species mean trait values along the climate gradient, and (ii) a within-species component, represented by the solid blue line, reflecting the average trait response within species across the climate gradient. Light blue lines show responses of individual taxa. The relative slopes of the among- and within-species components determine their contributions to the overall pattern (grey line). (c) Different combinations of among- and within-species contributions can generate similar overall trait–climate patterns. Components may contribute equally, act in the same direction (co-gradient), or oppose one another (counter-gradient). In some cases, the overall relationship is dominated by species turnover, whereas in others it is driven primarily by within-species adjustment.

**Table 1.** List of seven traits used in this study and their predicted trends with mean annual precipitation (MAP) and mean annual temperature (MAT). Hypothesised magnitude of within-species contributions to trait-climate relationships are also provided for each trait.

| Trait | Predicted response to MAP | Predicted response to MAT | Basis for trait–climate trends | Hypothesised within-species contribution to trait–climate trends |
|---|---|---|---|---|
| Leaf-internal to ambient CO<sub>2</sub> ratio<br>(C<sub>i</sub>:C<sub>a</sub>; unitless) | ↑ | ↑ | Lower C<sub>i</sub>:C<sub>a</sub> reflects greater CO<sub>2</sub> drawdown due to increased carboxylation capacity (for a given rate of stomatal conductance) at dry and cold sites (Wright et al. 2003, Prentice et al. 2014). | **High** – C<sub>i</sub>:C<sub>a</sub> is a physiological trait that can vary strongly within species due to independent adjustment of stomatal conductance and carboxylation capacity across environmental gradients. |
| Leaf N per unit area<br>(N<sub>area</sub>; g m<sup>-2</sup>) | ↓ | ↓ | N<sub>area</sub> substantially reflects investment in the carboxylating enzyme, Rubisco (Walker et al. 2014). Higher N<sub>area</sub> should therefore enhance carbon uptake relative to water loss in dry climates and mitigate temperature-related constraints on enzyme activity in cold climates (Wright et al. 2003, Prentice et al. 2014). | **Moderate to high** – N<sub>area</sub> integrates physiological investment with leaf structure; therefore, within-species adjustment is expected to be more constrained than purely physiological traits such as C<sub>i</sub>:C<sub>a</sub>. |
| Sapwood : leaf area ratio<br>(S<sub>A</sub>:L<sub>A</sub>; mm<sup>2</sup> cm<sup>-2</sup>) | ↓ | ↓ | Higher sapwood area relative to leaf area offsets reduced hydraulic conductivity linked to smaller vessels in dry sites and linked to small vessels that reduce freeze–thaw embolism risk, together with increased water viscosity, in cold sites (Brodribb and Feild 2000, Xu et al. 2026) | **Moderate** – allocation to stem and leaf tissues can both be changed within species in response to environmental conditions, allowing greater adjustment overall than is possible for the component traits individually. |
| Leaf mass per area<br>(LMA; g m<sup>-2</sup>) | ↓ | ↓ | Higher LMA is associated with longer leaf lifespans in dry and cold environments and contributes to higher N<sub>area</sub> under both conditions by increasing investment in photosynthetic compounds through thicker mesophyll tissues (Wright and Westoby 2002, Dong et al. 2022, Wang et al. 2023). | **Low to moderate** – reflects presumed stronger constraints on within-species variation in leaf morphology compared with physiological or nutrient traits. |
| Leaf area<br>(LA; mm<sup>2</sup>) | ↑ | ? | Smaller leaves have thinner insulating boundary layers, track air temperature more closely, and reduce the need for transpirational cooling in dry environments (Gates 1965, Wright et al. 2017). Small leaves are advantageous *both* at very hot and very cold sites (Wright et al. 2017). | **Low to moderate** – as above |
| Wood density<br>(WD; g cm<sup>-3</sup>) | ↓ | ? | Higher wood density is linked to a greater fibre wall fraction, which enhances drought tolerance by increasing resistance to xylem collapse in dry environments (Hacke et al. 2001, Zanne et al. 2010) | **Low** – reflecting its role as a structural trait associated with long-lived woody tissues, for which within-species adjustment across environmental gradients is likely to be less than that of leaf traits |
| Seed mass (mg) | ? | ? | Higher productivity environments (e.g. higher MAP) select for larger seed mass (Falster et al. 2008). Alternatively, larger seeds are required to tolerate harsh environments (e.g. high MAT, low MAP; Muller-Landau 2010) | **Moderate to high** – variation in maternal provisioning in response to local conditions in addition to local adaptation effects. |

## RESULTS

Across the seven traits considered, data at two or more sites were available for 1867 taxa, ranging from 84 unique taxa for S_A:_L_A t_o 1563 species for LMA (Extended Data Fig. 1). Traits varied widely with ranges comparable to those in global datasets (e.g., Díaz et al. 2016), especially seed mass (> six orders of magnitude variation across taxon-cell means, 0.1% to 99.99% quantile), leaf area (> five orders of magnitude variation), and LMA (77-fold variation; Table S1). C_i:_C_a v_aried the least, varying just 2.2-fold across all observations. Variance decomposition showed that the proportion of trait variance attributable to among-versus within-species (i.e., residual) variation differed between traits and functional groups (Extended Data Table 1). Across all species, seed mass exhibited the lowest proportion of within-species variation (0.06, meaning that 94% of variance was among species), followed by leaf area (0.08), LMA (0.15), N_area (_0.22), wood density (0.29), and C_i:_C_a (_0.36). S_A:_L_A_ showed the highest proportion (0.45), indicating nearly equal within- and among-species variation. Across all traits, within-species variation accounted for a greater proportion of total variation in non-woody than woody taxa (Extended Data Table 1), particularly for C_i:_C_a (_0.81 vs. 0.35) and leaf area (0.23 vs. 0.06).

### Relative contributions of turnover and within-species variation to trait–climate relationships

Given these differences in the total amount of within- and among-species trait variation, we next asked how much of the variation in each component was associated with climate. Climate varied widely across the dataset, from 120 to 7625 mm MAP and 4.0 to 29.0 °*C* MAT. We quantified the relative magnitude of within- and among-species (turnover) contributions to overall trait–climate relationships (*S_w_*, see Methods). Values of *S_w_* approaching 1 indicate stronger within-species contributions, while values approaching 0 indicate stronger turnover contributions.

When functional groups (woody, non-woody) were combined, among-species variation was the dominant component of trait variation associated with MAP, with all median *S_w_* values being below 0.5 (Fig. 2). This indicated that more than half of each trait–MAP relationship was attributable to among-species variation. Among-species contributions were strongest for leaf area, seed mass and wood density (Table S2), indicating that relationships between these traits and rainfall were driven primarily by species turnover rather than within-species trait changes. Relative within-species contributions increased progressively from LMA to C_i:_C_a, t_o N_area._ S_A:_L_A h_ad a similar *S_w_* to C_i:_C_a a_nd N_area b_ut had a much wider posterior distribution than these traits.

**Figure 2.**
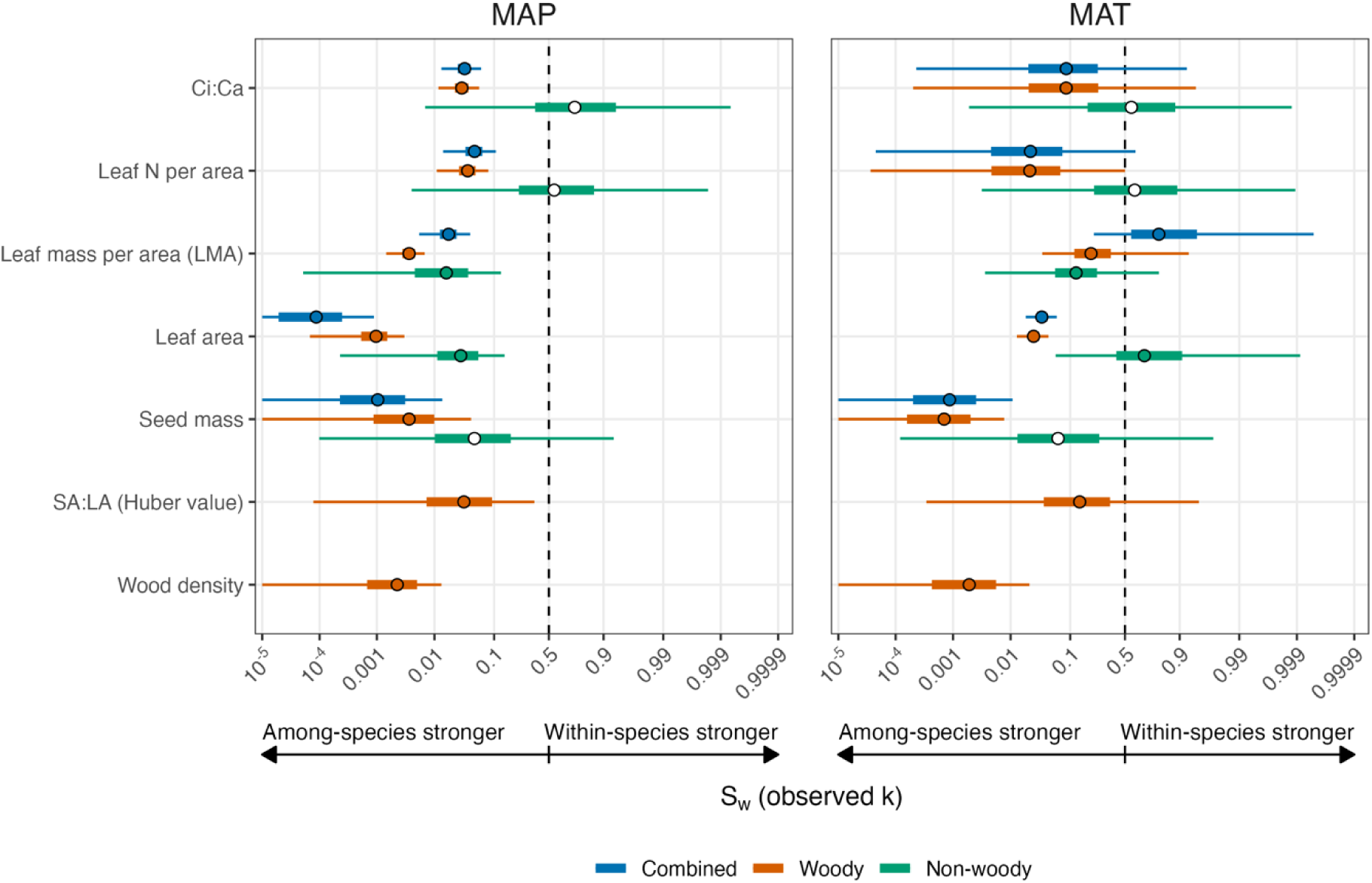
Within-species share of climate-associated trait variation *S_w_* along MAP and MAT gradients, evaluated at *κ* = *obs*. *S_w_* is the fraction of the trait variation generated by the fitted model along each gradient that arises from within-species adjustment rather than among-species responses, adjusted by the observed *κ* which is the climatic variation spanned within taxa relative to that spanned by taxon centroids. Values of *S_w_* below 0.5 indicate that among-species variation generates more of the climate-associated variation than within-species adjustment. Points show posterior medians, thick bars 50% and thin lines 95% credible intervals, on a logit scale; intervals extending beyond the axis limits are truncated at the panel edge. Points representing medians are unfilled when the 90% credible intervals of both among-(*B_m_*) and within-species (*B_w_*) contributions overlapped zero, indicating that neither component was significantly related to a given climate variable. Medians and 95% credible intervals are available in Table S2.

Trait–MAT relationships spanned a broader range of relative contributions (i.e., *S_w_*), although uncertainty around these estimates were greater than for MAP (Fig. 2, Table S2). Nevertheless, among-species contributions dominated relationships between MAT and seed mass, wood density and, to a lesser extent, leaf area. In contrast, LMA was the only trait for which the median within-species contribution exceeded 0.5, indicating that more than half of the observed LMA–MAT relationship arose from within-species variation. For N_area,_ C_i:_C_a a_nd S_A:_L_A,_ credible intervals overlapped 0.5, indicating broadly comparable contributions from among- and within-species variation, but with considerable uncertainty around these estimates.

Considering functional groups separately, non-woody taxa tended to show greater within-species contributions than woody taxa across most traits–climate relationships, although these differences were generally not significant (based on overlap in the credible intervals of posterior distributions of the orange and green bars; Fig. 2; Table S2). Median *S_w_* values were generally below 0.5 in both woody and non-woody taxa, indicating that among-species variation was the dominant component driving trait responses to climate, irrespective of functional type. Functional group differences were most apparent for leaf area, particularly for MAT, where the dominant contribution shifted from among-species variation in woody taxa to within-species variation in non-woody taxa (based on the median). For N_area,_ seed mass and C_i:_C_a i_n non-woody taxa, the broad posterior distributions indicated greater uncertainty in the relative contributions of within- and among-species variation to their climate relationships.

### Directionality of among- and within-species contributions to trait–climate relationships

The relative contributions described above indicate the relative importance of within-versus among-species variation to a given trait–climate relationship, but do not identify the direction or strength of their relationships with climate. We therefore examined the underlying within- and among-species relationships separately (Figs 3, 4) and compared them with the overall relationship and predictions in Table 1.

**Figure 3.**
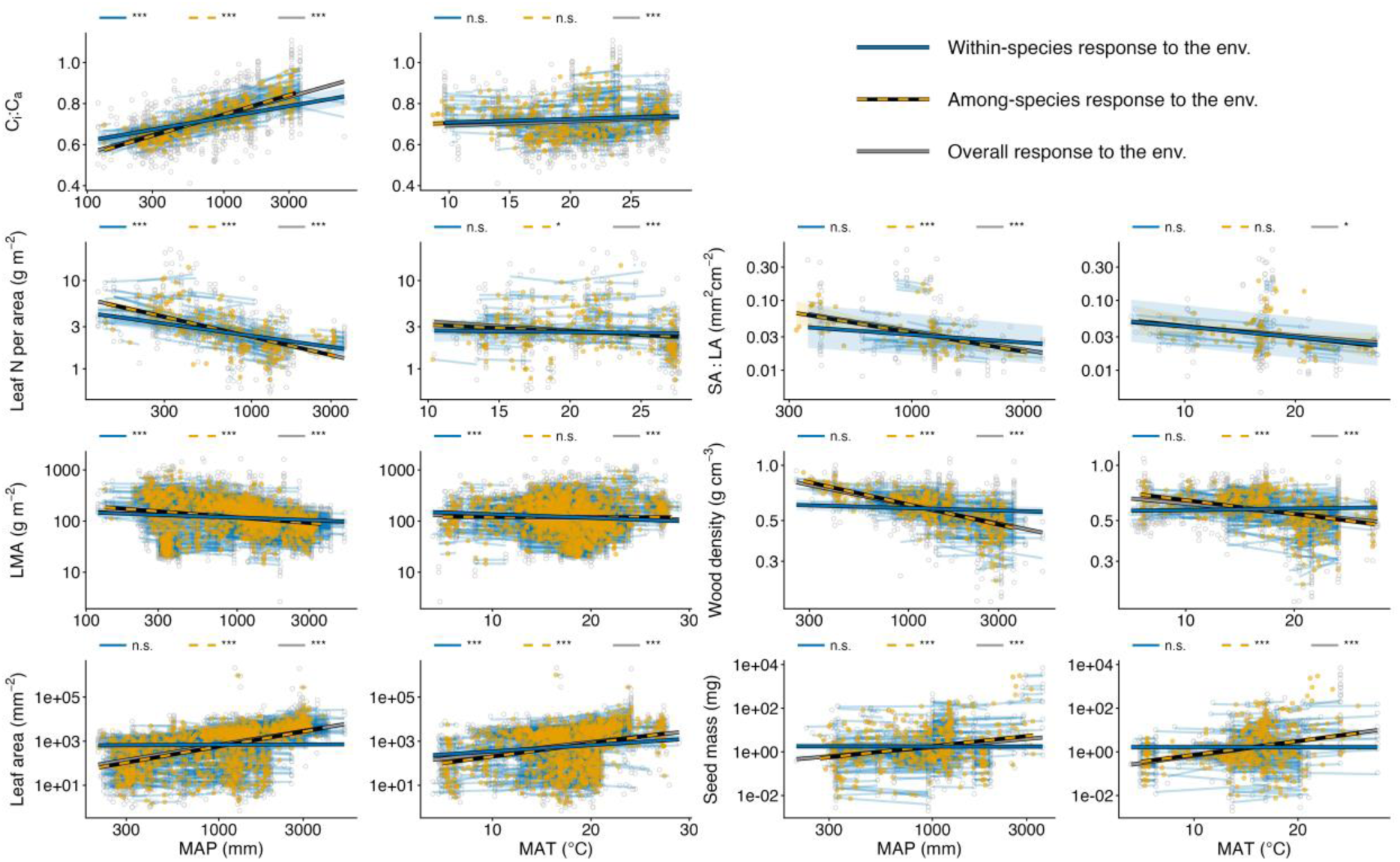
Decomposition of the relationship between traits and mean annual precipitation (MAP; left column) and mean annual temperature (MAT; right column) into among-species (yellow; dashed lines) and within-species components (blue; solid lines). The grey, unfilled points are trait observations (taxon means per grid cell) and the orange points are taxon-specific predicted trait values (on the y-axis) arranged by climate centroid (on the x-axis), being the mean MAP or MAT across all trait observations of each taxon. The grey line is fitted from an ordinary least squares regression of the raw observations against MAP or MAT. The fitted orange line is the estimated response of trait observations to variation in the MAP or MAT centroid (i.e.*B_m_* ⍰ in equation 4). Each semi-transparent blue line is the estimated taxon-specific response to deviation from its climate centroid and the solid blue line is the global estimate of the effect of deviation (i.e.*B_w_* in equation 4), centred at the median taxon centroid, where it intersects the across-species line. The fitted lines for the S_A:_L_A p_lot used parameters from the model fitted using MCMC sampling. The significance of parameter estimates is indicated by the legend at the top of each subpanel where n.s. is non-significant, * indicates p < 0.05, ** indicates p < 0.01 and *** indicates p < 0.001. Parameter estimates are reported in Table S3, S4.

Considering the combined effects of among-species (species turnover) and within-species variation (i.e., their overall trend; grey line, Fig. 3, Table S3), all traits were significantly related to MAP in the predicted directions (Table 1): species in wetter sites had, on average, lower N_area,_ LMA, wood density and S_A:_L_A,_ and higher C_i:_C_a,_ leaf area and seed mass. Among-species slopes were significant for every trait (yellow dashed lines, Fig. 3, left column; Table S4) and matched the direction of the overall trend, driving the dominant among-species contributions reported above. The fitted within-species trait–MAP slopes (bold blue lines, Fig. 3) were generally weaker, but were significant and in the same direction as turnover for N_area,_ LMA and C_i:_C_a._ Thus, within species sampled across sites, N_area a_nd LMA tended to decline with increasing MAP, while C_i:_C_a t_ended to increase. By contrast, wood density, S_A:_L_A,_ seed mass and leaf area showed no significant within-species association with MAP, indicating that their overall MAP relationships arose primarily through species turnover.

Overall trait–MAT trends were generally weaker than trait–MAP relationships but were significant for all traits and, where predicted, in the expected direction (Table 1): N_area,_ LMA, S_A:_L_A a_nd wood density responded negatively to MAT, while leaf area, seed mass and C_i:_C_a r_esponded positively (grey line, Fig. 3, right column; Table S3). Significant or close to significant (i.e. p < 0.10, C_i:_C_a p_ = 0.06; S_A:_L_A =_ 0.053) among-species slopes were observed for six of the seven traits (yellow dashed lines, Fig. 3, right column; Table S4), which generally matched the direction of the overall MAT trend. LMA was the exception, showing a non-significant among-species relationship with MAT but a significant within-species relationship, with individuals of the same species tending to have lower LMA in warmer sites. Leaf area was the only other trait with a significant within-species slope, increasing with MAT within species. For N_area,_ wood density, S_A:_L_A,_ seed mass and C_i:_C_a,_ the average within-species relationships with MAT were non-significant, indicating weak or inconsistent within-species contributions to the overall MAT trends. Shallower trait–MAT trends, especially for the within-species component, explained the wider posterior distributions in Figure 2 for MAT relative to MAP.

### Functional group comparison of among- and within-contributions to trait–climate relationships

Differences in *S_w_* between functional groups emerged primarily because woody taxa exhibited stronger among-species slopes in response to climate, rather than differences in the strength of the within-species slope (Fig. 4). Among-species (turnover) slopes differed significantly between functional groups (woody and non-woody, orange and green dashed lines respectively, Fig. 4, Extended Data Table 2) for all trait-MAP relationships except N_area,_ which declined similarly with MAP in both groups (Fig. 4, conditional effect for non-woody taxa non-significant [Table S5]). For C_i:_C_a,_ seed mass and leaf area, woody taxa exhibited steeper turnover slopes, indicating stronger among-species increases in these traits with increasing MAP than in non-woody taxa. LMA showed the strongest contrast between functional groups, with turnover-MAP slopes occurring in opposite directions: among-species LMA increased significantly with MAP in non-woody taxa but decreased significantly in woody taxa (Fig. 4; Table S5). Within-species trait responses to MAP were largely conserved across functional groups in a manner consistent with when they were combined. Leaf area was the exception, increasing with MAP very slightly within woody taxa but decreasing with MAP within non-woody taxa.

**Figure 4.**
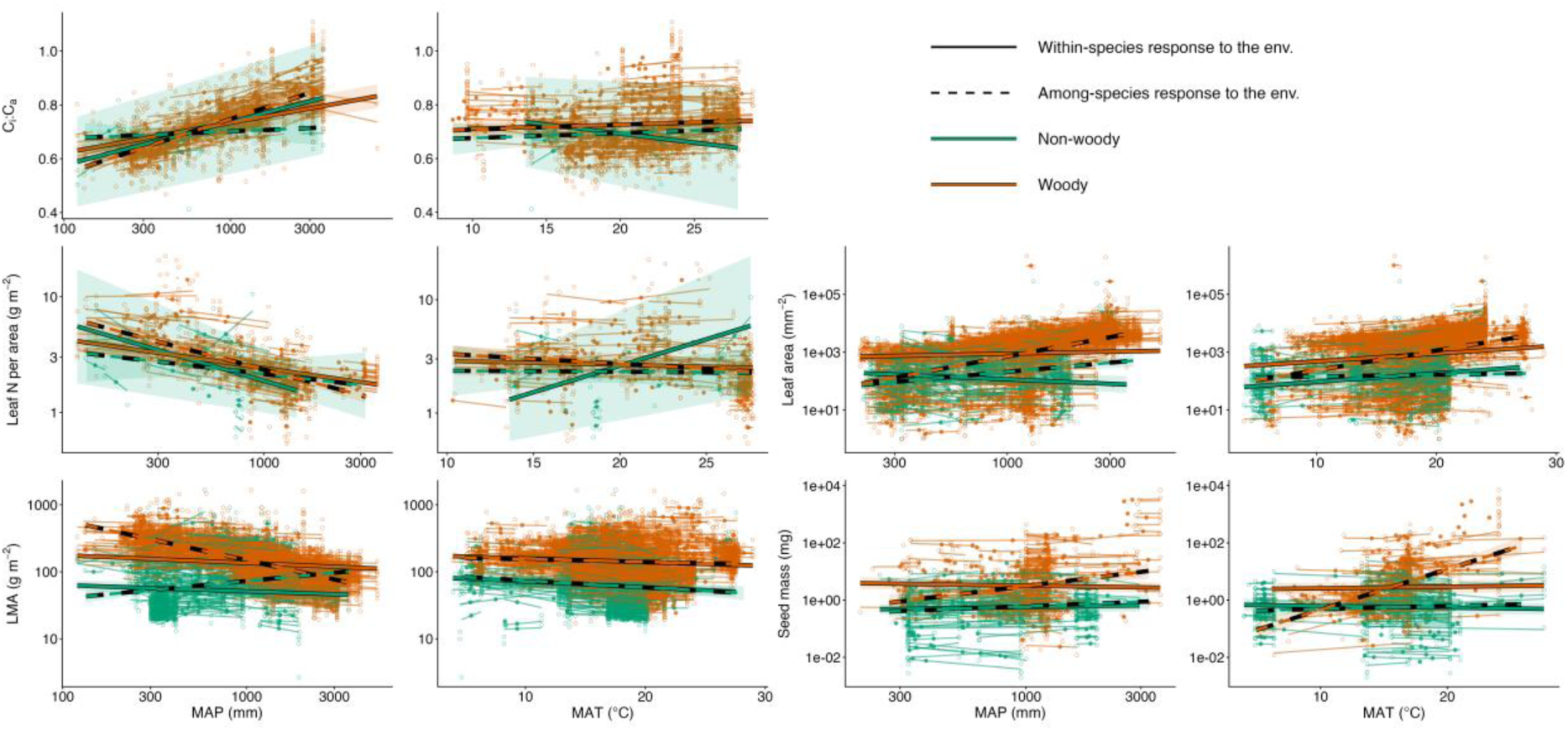
Decomposition of the relationship between traits and mean annual precipitation (MAP; left column) and mean annual temperature (MAT; right column) for woody (brown) and non-woody (green) taxa into among-(dashed line) and within-species (solid line) components. Comparisons between functional groups were limited to C_i:_C_a,_ N_area,_ LMA, leaf area and seed mass because stem traits (wood density and S_A:_L_A)_ are absent in herbaceous species. The coloured, unfilled points are trait observations (taxon means per grid cell) and the solid points are taxon-specific predicted trait values (on the y-axis) arranged by climate centroid (on the x-axis), being the mean MAP or MAT across all trait observations of each taxon. The fitted dashed line is the estimated response of trait observations to variation in the MAP or MAT centroid (i.e. in Equation 5). Each semi-transparent coloured line is the estimated taxon-specific response to deviation from its climate centroid and the solid line is the global estimate of the effect of deviation (i.e. in Equation 5), centred at the median taxon centroid, where it intersects the across-species line, extended across the full climatic range. Parameter estimates and significance of estimates are reported in Extended Data Table 2.

Differences between functional groups were less pronounced for MAT than for MAP (Fig. 4, right column). Nevertheless, turnover responses were generally stronger in woody taxa: significantly steeper positive slopes for leaf area and seed mass were observed in woody than non-woody taxa; non-woody turnover slopes were insignificant in both cases (Extended Data Table 2). Similarly, significant or close to significant turnover responses for N_area (_negative) and C_i:_C_a (_positive) were observed only in woody taxa, although these responses did not differ significantly between functional groups owing to substantial variation in the predicted slope estimates for the non-woody group (Table S4). By contrast, LMA showed similar negative turnover responses to MAT in both groups, with the woody turnover slope being slightly shallower. Within-species responses to MAT did not differ significantly between functional groups (Extended Data Table 2). Instead, consistent with patterns across all taxa, leaf area increased and LMA decreased similarly with MAT within both woody and non-woody taxa, and no other trait showed a significant within-species response to MAT in either functional group (Table S5).

## DISCUSSION

Drawing on a continental-scale trait dataset spanning 1867 plant species, this study examined how species turnover and within-species trait adjustment jointly shape trait–climate relationships along precipitation and temperature gradients. We find:

1. Species turnover was the dominant process shaping climate-driven trait variation, with within-species adjustment playing a secondary role.
2. The relative importance of turnover versus within-species adjustment differed markedly among traits, although this division was clearer along precipitation gradients than temperature gradients. Within-species adjustment contributed most strongly to climate-related variation in leaf physiological and biomass allocation traits, and least to leaf area, seed mass and wood density; LMA depended on the climate variable considered.
3. Species turnover slopes were generally steeper and contributed more strongly to trait–climate relationships in woody taxa, whereas within-species contributions were generally stronger in non-woody taxa.

The dominance of species turnover across all seven traits suggests that broad-scale trait–climate relationships have historically been shaped principally by changes in species composition rather than by trait adjustment within species. This key finding provides insight into how vegetation may track future changes in climate. In principle, as climatic conditions shift, so too should the ‘optimal’ trait value associated with a given location. To the extent that these spatial patterns provide a basis for future temporal responses, our results indicate that plant traits will track changing climatic conditions primarily through the replacement of species with different trait values rather than through large *in situ* adjustments within existing species. Future changes in the functional composition of plant communities may therefore be constrained by the relatively slow process of species dispersal and replacement, potentially causing trait shifts to lag behind climatic changes (e.g., Quetin et al. 2023).

However, many species can persist well beyond their historical geographic distributions, e.g., in revegetation sites and botanic gardens (Laughlin and McGill 2024). Such expansions of climatic niche space may be achieved through trait changes; for example, North American pines transplanted to the Southern Hemisphere have shifted toward more drought-tolerant trait values (Fajardo et al. 2026). These observations imply greater capacity for within-species trait adjustment than may be inferred from spatial trait–climate associations such as those examined here. One potential explanation is that gene flow among populations occupying different environments constrains local differentiation, resulting in more similar trait values across populations in different climates. This could generate shallower within-species trait–climate relationships than might otherwise be observed between genetically isolated populations of the same species (Kirkpatrick and Barton 1997, Yang et al. 2026). Thus, the relatively small within-species contributions observed in this study may underestimate the capacity of species to persist *in situ* under future climate change through trait adjustment. Even so, our analyses reveal that traits differ in the extent of within-species adjustment, suggesting that some aspects of plant form and function may respond more readily to climatic variation than others.

In applying a common analytical framework across a broad suite of traits we found general support for our hypothesised rank order of within-species contributions to trait–climate relationships (Table 1). The rank order of traits was most clearly expressed with respect to precipitation but was less clear for temperature owing to generally weaker overall trait-temperature relationships, as reported in previous continental-scale trait–climate analyses from Australia and North America (Towers et al. 2023, Britton et al. 2026, Vinod et al. 2026), albeit not globally (Moles et al. 2014).

The strongest within-species contributions to climate-related variation were detected for traits associated with physiological function and biomass allocation, namely C_i:_C_a,_ N_area (_less clearly for MAT than MAP) and S_A:_L_A._ These strong within-species contributions are consistent with previous studies of leaf physiological and nutrient traits (Dong et al. 2020, Tian et al. 2024), and with our expectations given the capacity for these traits to adjust over relatively short timescales. C_i:_C_a r_eflects the balance between stomatal conductance and carboxylation, processes that can adjust over minutes to weeks or months (Odé et al. 2025); N_area i_s closely linked to photosynthetic capacity because a large proportion of leaf nitrogen is invested in the enzyme Rubisco (Walker et al. 2014, Dong et al. 2017), and can vary over seasonal timescales (Bloomfield et al. 2018). S_A:_L_A l_ikewise showed high within-species contributions across both climate gradients, consistent with previous evidence that it can shift across environmental gradients within a species (Mencuccini and Grace 1995, Rosas et al. 2019). Adjustment of leaf area through leaf shedding in response to climatic stress (Bucci et al. 2005, O’Grady et al. 2009) provides one mechanism through which S_A:_L_A c_an shift relatively readily within species, compared with changes to vessel dimensions occurring over longer timescales through the development of new woody tissue.

By contrast, seed mass, wood density and leaf area (for MAP) showed relatively small within-species contributions to overall trait–climate relationships. Such weak contributions could arise because a trait has (1) little inherent within-species variation, (2) within-species variation that is not systematically related to climate, or (3) climate-related variation that differs in direction among species and therefore produces a weak median response. Our results largely reflect the first two situations in these three traits.

For leaf area, we observed little inherent within-species variation overall (supporting point 1 above), while individual species’ responses to MAP were centred around zero, resulting in a non-significant overall within-species response (supporting point 2). This is consistent with previous studies reporting weak within-species leaf area responses across Australian climatic and geographical gradients (Dong et al. 2020, Guo et al. 2024). However, the steeper within-species response to MAT suggests that even limited inherent variation can be systematically patterned with some aspects of climate but not others, here with MAT but not MAP. Seed mass showed little inherent within-species variation (6%), and within-species responses to both MAP and MAT were tightly clustered around zero (Fig. S2), supporting both points 1 and 2. The former may partly reflect an evolutionary constraint imposed by the seed size–number trade-off (Smith and Fretwell 1974), while the latter may arise because much of the seed mass variation that does occur is among seeds within individual maternal plants rather than being systematically patterned with climate (Michaels et al. 1988, Obeso 1993, Westoby et al. 1996).

Wood density showed a significant within-species response to MAP, although this was small relative to the among-species response, and no within-species contribution to its weak relationship with MAT. Thus, unlike leaf area and seed mass, within-species variation in wood density was systematically patterned with precipitation but contributed relatively little to the overall relationship. Within-species increases in wood density under drier conditions have been reported at regional- and global-scale (Anderegg et al. 2021, Fischer et al. 2026), potentially enhancing drought resistance through increased fibre wall fraction (Hacke et al. 2001). The absence of a comparable MAT response to that observed in global analyses (Fischer et al. 2026) may reflect differences in the climatic ranges represented between studies, with global analyses encompassing environments with colder winters where freeze-thaw cycles could intensify selection for embolism resistance via narrow vessel diameters and denser wood (Olson et al. 2020).

One notable exception to our hypothesised ranking of traits was the relatively high within-species contribution to LMA variation across both climate variables, particularly for MAT. Increasing MAT was associated with lower LMA through within-species adjustment, with no detectable corresponding among-species (turnover) response. In contrast, the high contribution for MAP was partly inflated by opposing turnover responses in woody and non-woody taxa, which cancelled when pooled and produced a shallow overall turnover slope (Fig. 4). Analysed separately, LMA was intermediate between the physiological and biomass allocation traits, and leaf area, seed mass and wood density, broadly aligning with our expectations and conclusions from Dong et al. (2020). Thus, despite some differences between climate gradients, our results generally supported our hypothesised ranking, with the greatest within-species contributions generally occurring in physiological and allocation traits and the lowest in leaf area, wood density and seed mass.

Trait–climate relationships were generally stronger in woody than in non-woody species, consistent with previous studies (Šímová et al. 2018, Towers et al. 2023). Our results show that this pattern arises primarily through species turnover: woody taxa exhibited steeper turnover responses to climate, while within-species contributions were broadly similar between functional groups, or contributed proportionally more to non-woody taxa. In other words, a given change in climate was associated with larger trait shifts in woody than non-woody species, primarily because their turnover responses were stronger rather than because they showed greater within-species adjustment. This pattern is consistent with the idea that woody plants experience macroclimate more directly (Bailey and Sinnott 1916), while many non-woody species are buffered by occurring in favourable microclimates, such as beneath canopies (De Frenne et al. 2019) or by avoiding unfavourable periods through seed-bank persistence (Dwyer and Erickson 2016). Together with their generally greater phenotypic plasticity and shorter generation times (Thomson et al. 2011), this suggests that non-woody taxa may rely less on species turnover in response to future climate change. Differences in the degree of compositional change between woody and non-woody taxa may generate novel species combinations and thus interactions among co-occurring species. Understanding how these dynamics scale up to influence ecosystem function will be an important avenue for future research.

Using a continental-scale plant trait dataset, we show that climate-related trait variation is shaped principally by species turnover, with within-species trait adjustment playing a secondary role. Applying a common analytical framework across a broad suite of traits revealed substantial differences among traits in their capacity for within-species adjustment, providing a general ranking of the capacity for different aspects of plant function to respond to climatic variation within species. We also demonstrate that stronger trait-climate relationships in woody taxa arise primarily through steeper turnover responses to climate, while within-species contributions are broadly similar between woody and non-woody taxa. These findings reveal which aspects of plant function are most likely to persist through within-species adjustment, and which will depend on species replacement, as climates change.

## METHODS

### Trait data

AusTraits (Version 7.0.0; Falster et al. 2021) is a harmonised database of plant functional traits representing ∼5000 taxa for the traits of interest in this study and ∼1900 taxa with observations in two or more distinct locations. We selected seven functional traits to analyse according to a number of criteria, being that they: 1) were sampled broadly across focal climate gradients, 2) captured different aspects of plant ecophysiological strategy, and 3) potentially represent a range of relative magnitudes of within-species variation compared to among-species (turnover) variation across climatic gradients (Table 1). For example, while structural traits such as leaf area may exhibit relatively limited within-species variation, leaf chemical traits and water-use efficiency are often found to be relatively plastic, allowing for acclimation in different environments (Siefert et al. 2015, Dong et al. 2020)

After selecting the focal traits, we applied the following filters to generate a suitable subset for this study. Firstly, we used the “taxon_rank” classification to remove all trait records that were classified at an organisational level higher than “species” (i.e. genus, family, e.g. “*Eucalyptus* sp.”). Secondly, to ensure that observed trait variation represented “natural” variation, we removed data sourced from manipulative experiments, although we retained data from two studies where it was noted that data were from naturally occurring, unmanipulated samples (in both cases control plots). Similarly, we also removed data obtained from laboratory, glasshouse and arboretum populations. Thirdly, we removed data that were recorded as species-level averages or were sourced from herbarium records or expert opinion – as these have no site location for which climate data can be sourced. Finally, we only retained data with associated geospatial information. We also applied additional filters to certain traits. For leaf area of compound-leaves species, we only retained observations where they described the size of leaflets rather than for entire leaves (Towers et al. 2023). For leaf *δ*^13^*C*, we filtered the dataset to exclusively include C3 taxa, i.e. removing species listed in AusTraits as having C4 or CAM photosynthesis. If the photosynthetic type of a given taxa was unknown, we then removed observations where leaf *δ*^13^*C* was less negative than −20‰, as this most likely indicates species C4 or CAM rather than C3 species (Crayn et al. 2015). We also validated this cut-off by visually inspecting histograms grouped by known photosynthetic pathway.

Leaf *δ*^13^*C* data were converted to the ratio of the intracellular carbon dioxide concentration to atmospheric carbon dioxide concentration (i.e. *C_i_*: *C_a_*) by first correcting for the isotopic signature of air (Cernusak and Ubierna 2022):

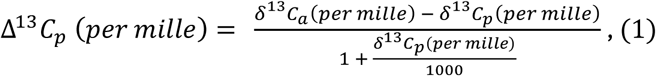

where Δ^13^*C* is the corrected value, *δ*^13^*C* is the carbon isotopic signature of the plant sample and *δ*^13^*C* is the carbon isotopic signature of air.

To account for variation in *δ*^13^*C_a_* through time, we estimated *δ*^13^*C_a_* based on the year of collection (CE) of each observation of *δ*^13^*C_p_* according to the following empirical equation:

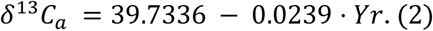

*C_i_*: *C_a_* was then calculated according to the following equation:

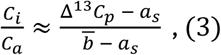

where *a_s_* is 4.4 (per mille) and *b* is 27 (per mille).

After these filters and transformations were applied, the Australian continent was divided into grid cells at 30 arc-second resolution (approximately 1 km at the equator), to be consistent with climate data (described below). For each trait, we then filtered the dataset down to taxa with trait observations in two or more grid cells. After log-transforming all traits but C_i:_C_a,_ we then calculated the mean value per cell of a given trait for a given taxon, including across datasets, to account for potential bias emerging from oversampling of traits in certain locations. We created a plant growth form variable with two states: woody and non-woody, by combining groups from a more specific plant growth form grouping available for almost all taxa in AusTraits. To do this, we used a binary classification system to assign all taxa with the following keywords: “herb”, “fern”, “tussock”, “geophyte”, “lycophyte”, “basal_large”, “graminoid” and “hummock” to be non-woody while all remaining taxa with growth form data available were assigned to be woody. Occasionally, taxa (43 in total) had multiple keywords which could be considered either non-woody or woody (e.g. “herb shrub”) and in these cases we assigned them to be non-woody – these taxa presumably express multiple life history strategies including annual above-ground biomass but perennial belowground structures, conferring the “avoider” strategies consistent with other non-woody taxa (e.g. *Vittadinia cuneata*).

### Climate data

Climate data were obtained from the standard set of long-term (1970-2020) annual bioclimatic variables from WorldClim (version 2.1; Fick and Hijmans 2017). The variables that we selected were mean annual precipitation (MAP) and mean annual temperature (MAT), using the highest spatial resolution available (30 arc-seconds [approximately 1 km at the equator]). MAP and MAT were pragmatically selected as variables which 1) represent two key dimensions of the environment being moisture availability and temperature, 2) are widely considered to structure global vegetation patterns and associated variation in plant functional traits (Schimper 1902, Warming 1909, MacArthur 1984, Fonseca et al. 2000, Moles et al. 2014) and 3) are easily interpretable and comparable to other studies.

### Statistical analyses

To analyse the data, we employed a recently developed random intercepts and slopes linear modelling approach which partitions among- and within-species changes in traits with climate by simultaneously regressing a trait against species-specific climate centroids and the deviation of a given observation of a species’ trait, in climate space, from its own centroid (Fischer et al. 2026). The component about centroids captures community-wide trends emerging from species replacement while the deviations represent the response of within-species variation to climate, which can vary based on species. This approach differs from earlier trait-gradient analyses which considered how strongly individual species respond to the environment relative to community-wide trait means (Ackerly and Cornwell 2007, Lepš et al. 2011, Dong et al. 2020, Tian et al. 2024). Two advantages of the approach utilised here are that it does not depend on exhaustive sampling of traits in each location and therefore facilitates the use of harmonised trait databases such as AusTraits, and that it uses the abiotic environment (here MAP, MAT) as predictors, rather than trait means themselves.

Taxon-level climate centroids were calculated separately for each climate variable. Prior to calculating centroids, MAP was log-transformed, and then both MAP and MAT were mean-centred and scaled by 1 standard deviation. Then, for each taxon, the mean value of the scaled climate data associated with all observations of the seven traits (if available) for that taxon was calculated. We elected to calculate centroids based on taxon occurrences in AusTraits, as opposed to other databases such as GBIF (GBIF 2026). While the latter may yield a more accurate representation of the true centre of a species’ distribution, the centroid in the former case represents the modal climate within the geographic distribution of the observed trait data (Fischer et al. 2026). To account for bias potentially emerging from highly concentrated sampling of traits in certain locations, each grid cell contributed only once to each trait-taxon centroid. Deviations from the centroid were then calculated by subtracting the taxon-specific centroid from the scaled value of the environmental variable associated with each trait observation (Fischer et al. 2026).

Again, to account for uneven sampling of traits of a given taxa across a gradient, we calculated the trait-taxon mean per grid cell. Consequently, the level of observation is the grid cell trait-taxon average. Prior to modelling the response of traits to climate, we first assessed the relative proportion of variation for each trait that is attributable to within-species variation, whether or not it was related to climate. To do this, we fit an intercept-only mixed effects model of each trait with taxon identity as a random intercept. The residual variation as a fraction of the residual variation plus the between-taxa variation was then taken as an indicator of within-species variation for each trait, with values closer to 0 indicating a lower proportion of within-species variation and values closer to 1 indicating a higher proportion of within-species variation.

Geographic variation in each trait was then modelled as a function of the centroid and centroid deviations in relation to climatic predictors:

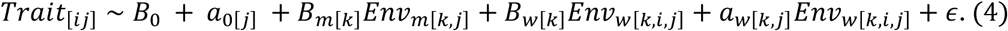

Here, observations, *Trait*_[*ij*]_, of a given trait across all individuals, i, and species j, were fit as a function of *Env_m_*_[*k*,*j*]_, being the centroid of the environmental variable, k (i.e. either MAP or MAT; fit separately) for species j across all observations for that species, and *Env_w_*_[*k*,*i*,*j*]_, being the deviation between the environmental variable for a given observation and *Env_m_*_[*k*,*j*]_. Overall, the model quantifies the effect of among-species variation along environmental gradients owing to species turnover (i.e. *Env_m_*_[*k*,*j*]_) and within-species variation emerging from species varying their traits as they move away from their typical environment (i.e. *Env_w_*_[*k*,*i*,*j*]_) on emergent trait values. *B*_0_ is a global intercept, representing the average value of a given trait and *B_m_*_[*k*]_ is a fixed effect parameter representing the effect of a unit change in *Env_m_*_[*k*,*j*]_ on species centroids. *a*_0[*j*]_and *a_c_*_[*k*,*j*]_are the random intercept parameter and the random slope parameter for the effect of *Env_w_*_[*k*,*i*,*j*]_ on a given trait, both of which vary by species. Importantly, this means that a species’ response to *Env_w_*_[*k*,*i*,*j*]_ can vary in both magnitude and sign. Moreover, as we did not have an *a priori* hypothesis that a species’ average trait value is related to the magnitude and sign of its within-species response to the environment, we did not assume a correlation between *a*_0[*j*]_ and *a_w_*_[*k*,*j*]_. *B_w_*_[*k*]_ is a fixed effect parameter representing the average effect of *Env_w_*_[*k*,*i*,*j*]_, being the average within-species trait response to a unit deviation from the environmental centroid. Finally, *ε* represents residual error.

We also explored whether the among-and within-species responses to the environment depended on plant growth form. To do this, we added plant growth form (PGF) as an interacting variable to the above model:

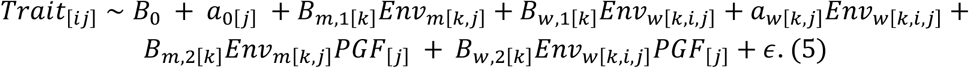

Because the degree of variation in species-level intercepts and slopes may vary between woody and non-woody taxa, we separately fitted the variance term of the random intercept and slope for each functional group, and compared this model set to a simpler model set with a pooled variance term based on AIC. For all trait and environmental variable combinations, separate variance terms either improved model fit (i.e. AIC > 2; Table S6) or were equivalent, so we retained separate variance terms.

We used the glmmTMB function from the glmmTMB package (Brooks et al. 2017) to fit the models using maximum likelihood. To ensure model convergence, we fit traits to MAP and MAT separately.

Finally, to evaluate the relative magnitude of within- and among-species variation, we calculated the fraction of climate-associated trait variation along each climate gradient contributed by each component as:

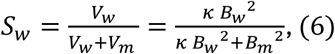

where *V_w_* and *V_m_* are the variances in fitted trait values that each component generates across the observations for within- and among-species components, respectively, given by

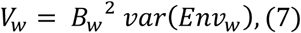

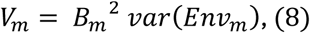

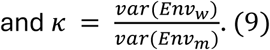

As *Env_w_*_[*k*,*i*,*j*]_ is a deviation from each taxon’s climate centroid, the two predictors are close to uncorrelated by construction (|r| < 0.15 for all traits; Table S2), so the two variances can be treated as additive. Both variances are taken across all observations in the fitted dataset (the taxon × grid-cell records).

*S_w_*varies from 0 to 1, with values closer to 1 indicating greater contribution from within-species trait responses and values closer to 0 indicating a greater contribution of the among-species trait response to the observed climate response.

Because *κ* is calculated from the locations at which traits were measured rather than from the full distribution of each taxon, the observed value, *k_obs_*, is likely to underestimate the true value. Indeed, trait records sample only part of each taxon’s climatic range and centroids estimated from few records are imprecise, which inflates var(*Env_m_*_[*k*]_). We therefore treated *S_w_* evaluated at *κ_obs_* as a lower estimate of the variance explained by within-species variation. As an upper estimate, we set *κ* = 1, equivalent to assuming that each taxon spans internally as much climatic variation as is spanned across all taxon centroids, or that half of all climatic variation in the dataset occurs within taxa (Fig. S4). Since *S_w_* increases monotonically with *κ*, we also report *S_w_* across the interval *k_obs_* < *k* < 1 (Fig. S5).

Primarily, we were concerned with whether the credible interval (i.e. the 2.5^th^ and 97.5^th^ quantile) of *S_w_*(*κ*) overlapped 0.5, indicating that each component did not contribute significantly more strongly to climate-associated variance, and whether the credible intervals of the joint distribution in response to a given climate variable overlapped between traits, indicating differences between traits in the relative contribution of components. To generate credible intervals, we refit the models above using MCMC sampling via the brm function in the brms R package (Bürkner 2017). We then extracted the posterior distributions to calculate the differences described above, generating a joint posterior distribution for each. Bayesian models were fit using 4 chains with 7,500 iterations each. For leaf area when all taxa were combined, it was necessary to fit the model using 15,000 iterations. Successful model convergence was assessed by ensuring that R-hat <= 1.05 for all parameters estimates. Maximum likelihood models failed to converge in some cases for S_A:_L_A a_nd so we exclusively report parameter estimates and visualisations emerging from MCMC sampling.

In order to ground within- and among-species trait responses to the environment in the context of overall trait-environment responses, we also conducted simple linear models of traits as a function of either MAP or MAT, including functional group as an additional analysis (grey lines in Fig. 3, Table S1, Table S8).

## DATA AVAILABITY STATEMENT

All data used in this study is available at https://zenodo.org/records/15718081. All code required to extract data and repeat analyses will be made publicly available upon article acceptance via a GitHub repository.

## AUTHOR CONTRIBUTIONS

TGB and IJW conceived the initial study, with IRT and DSF subsequently contributing to its development. IRT and DSF analysed the data. TGB wrote the first draft with contributions from IRT and LJW. All authors contributed to subsequent versions of the manuscript.

## ACKNOWLEDGEMENTS

We thank the many data contributors to the AusTraits plant trait database, whose data made this study possible. TGB and IJW were supported by an Australian Research Council (ARC) Centre of Excellence grant (CE200100015); IRT and DSF by an ARC Linkage Project (LP230100049) and Discovery Project (DP240100177) grant; LJW by an ARC Discovery Early Career Researcher Award (DE250100704).

## EXTENDED DATA

**Extended Data Table 1.** Proportion of total trait variance attributable to within-species variation (WSV; i.e. residual variation) for seven traits across all taxa combined and for woody and non-woody functional groups separately based on an intercept-only random effects model with taxon as a random intercept. The remaining variance (1 − proportion shown) represents among-species variation. Values were unavailable for wood density and SA:LA in non-woody taxa. The median range in climate spanned by each taxon and the total climatic range across centroids is reported.

|  |  |  |  | Median range in |  | Centroid range |  |
|---|---|---|---|---|---|---|---|
| **Group** | **ITV fraction** | **Taxa** | **Median cells per taxon** | **MAP (mm)** | **MAT (°C)** | **MAP (mm)** | **MAT (°C)** |
| **Ci:Ca** |  |  |  |  |  |  |  |
| Combined | 0.36 | 422 | 2.0 | 221 | 1.4 | 3,232 | 19.5 |
| Woody | 0.35 | 399 | 2.0 | 225 | 1.5 | 3,232 | 19.5 |
| Non-woody | 0.81 | 23 | 2.0 | 167 | 1.0 | 2,891 | 13.9 |
| **Leaf N per area** |  |  |  |  |  |  |  |
| Combined | 0.22 | 228 | 2.0 | 198 | 0.8 | 3,031 | 17.3 |
| Woody | 0.22 | 204 | 2.0 | 214 | 0.8 | 3,031 | 17.3 |
| Non-woody | 0.26 | 24 | 2.0 | 152 | 0.5 | 1,245 | 13.0 |
| **Leaf mass per area (LMA)** |  |  |  |  |  |  |  |
| Combined | 0.15 | 1,563 | 3.0 | 288 | 2.0 | 3,451 | 23.3 |
| Woody | 0.19 | 1,217 | 3.0 | 328 | 2.0 | 3,451 | 22.4 |
| Non-woody | 0.19 | 346 | 3.0 | 177 | 2.2 | 2,969 | 21.9 |
| **Leaf area** |  |  |  |  |  |  |  |
| Combined | 0.08 | 1,262 | 3.0 | 300 | 2.0 | 3,644 | 22.5 |
| Woody | 0.06 | 974 | 3.0 | 332 | 2.2 | 3,644 | 21.6 |
| Non-woody | 0.23 | 288 | 2.0 | 231 | 1.8 | 2,849 | 21.9 |
| **SA:LA (Huber value)** |  |  |  |  |  |  |  |
| Woody | 0.45 | 84 | 2.0 | 172 | 0.5 | 2,763 | 21.2 |
| **Wood density** |  |  |  |  |  |  |  |
| Woody | 0.29 | 505 | 3.0 | 373 | 2.1 | 3,305 | 21.6 |
| **Seed mass** |  |  |  |  |  |  |  |
| Combined | 0.06 | 448 | 2.0 | 234 | 1.5 | 2,967 | 20.9 |
| Woody | 0.05 | 278 | 2.0 | 224 | 1.0 | 2,955 | 19.9 |
| Non-woody | 0.13 | 170 | 2.0 | 315 | 2.5 | 1,916 | 20.9 |

**Extended Data Table 2.** Fixed effect outputs for the decomposition of the relationship between traits and mean annual precipitation (MAP; left column) and mean annual temperature (MAT; right column) into among-(dashed lines) and within-species components (solid lines) based on functional groups.

|  | MAP |  |  | MAT |  |  |
|---|---|---|---|---|---|---|
| Variable | Estimate | Std. error | p value | Estimate | Std. error | p value |
| **Ci:Ca** |  |  |  |  |  |  |
| Intercept | 0.701 | 0.014 | <0.001 | 0.692 | 0.012 | <0.001 |
| Across-species slope | 0.009 | 0.013 | 0.517 | 0.009 | 0.013 | 0.471 |
| Within-species slope | 0.053 | 0.042 | 0.212 | -0.033 | 0.068 | 0.630 |
| Woody | 0.041 | 0.014 | 0.003 | 0.030 | 0.013 | 0.026 |
| Across-species slope × woody | 0.058 | 0.014 | <0.001 | -0.000 | 0.014 | 0.997 |
| Within-species slope × woody | -0.016 | 0.043 | 0.710 | 0.040 | 0.068 | 0.556 |
| **Leaf N per area** |  |  |  |  |  |  |
| Intercept | 0.774 | 0.140 | <0.001 | 0.815 | 0.119 | <0.001 |
| Across-species slope | -0.152 | 0.141 | 0.281 | -0.010 | 0.128 | 0.936 |
| Within-species slope | -0.386 | 0.251 | 0.124 | 0.519 | 0.391 | 0.184 |
| Woody | 0.104 | 0.144 | 0.469 | 0.233 | 0.127 | 0.067 |
| Across-species slope × woody | -0.198 | 0.145 | 0.172 | -0.087 | 0.134 | 0.518 |
| Within-species slope × woody | 0.195 | 0.254 | 0.443 | -0.557 | 0.396 | 0.159 |
| **Leaf mass per area (LMA)** |  |  |  |  |  |  |
| Intercept | 4.273 | 0.051 | <0.001 | 4.120 | 0.044 | <0.001 |
| Across-species slope | 0.197 | 0.048 | <0.001 | -0.116 | 0.048 | 0.017 |
| Within-species slope | -0.070 | 0.049 | 0.152 | -0.095 | 0.040 | 0.018 |
| Woody | 0.764 | 0.052 | <0.001 | 0.847 | 0.048 | <0.001 |
| Across-species slope × woody | -0.651 | 0.050 | <0.001 | 0.073 | 0.052 | 0.163 |
| Within-species slope × woody | -0.018 | 0.052 | 0.726 | 0.032 | 0.043 | 0.455 |
| **Leaf area** |  |  |  |  |  |  |
| Intercept | 5.301 | 0.106 | <0.001 | 5.082 | 0.102 | <0.001 |
| Across-species slope | 0.504 | 0.100 | <0.001 | 0.068 | 0.091 | 0.453 |
| Within-species slope | -0.243 | 0.132 | 0.067 | 0.329 | 0.131 | 0.012 |
| Woody | 1.180 | 0.122 | <0.001 | 1.598 | 0.120 | <0.001 |
| Across-species slope × woody | 0.555 | 0.117 | <0.001 | 0.690 | 0.117 | <0.001 |
| Within-species slope × woody | 0.352 | 0.139 | 0.011 | -0.028 | 0.136 | 0.838 |
| **Seed mass** |  |  |  |  |  |  |
| Intercept | -0.505 | 0.137 | <0.001 | -0.522 | 0.147 | <0.001 |
| Across-species slope | 0.245 | 0.149 | 0.101 | 0.117 | 0.113 | 0.304 |
| Within-species slope | 0.094 | 0.115 | 0.413 | -0.066 | 0.084 | 0.435 |
| Woody | 1.582 | 0.181 | <0.001 | 2.272 | 0.196 | <0.001 |
| Across-species slope × woody | 0.536 | 0.242 | 0.027 | 1.452 | 0.209 | <0.001 |
| Within-species slope × woody | -0.189 | 0.152 | 0.214 | 0.125 | 0.131 | 0.340 |

**Extended Data Figure 1.**
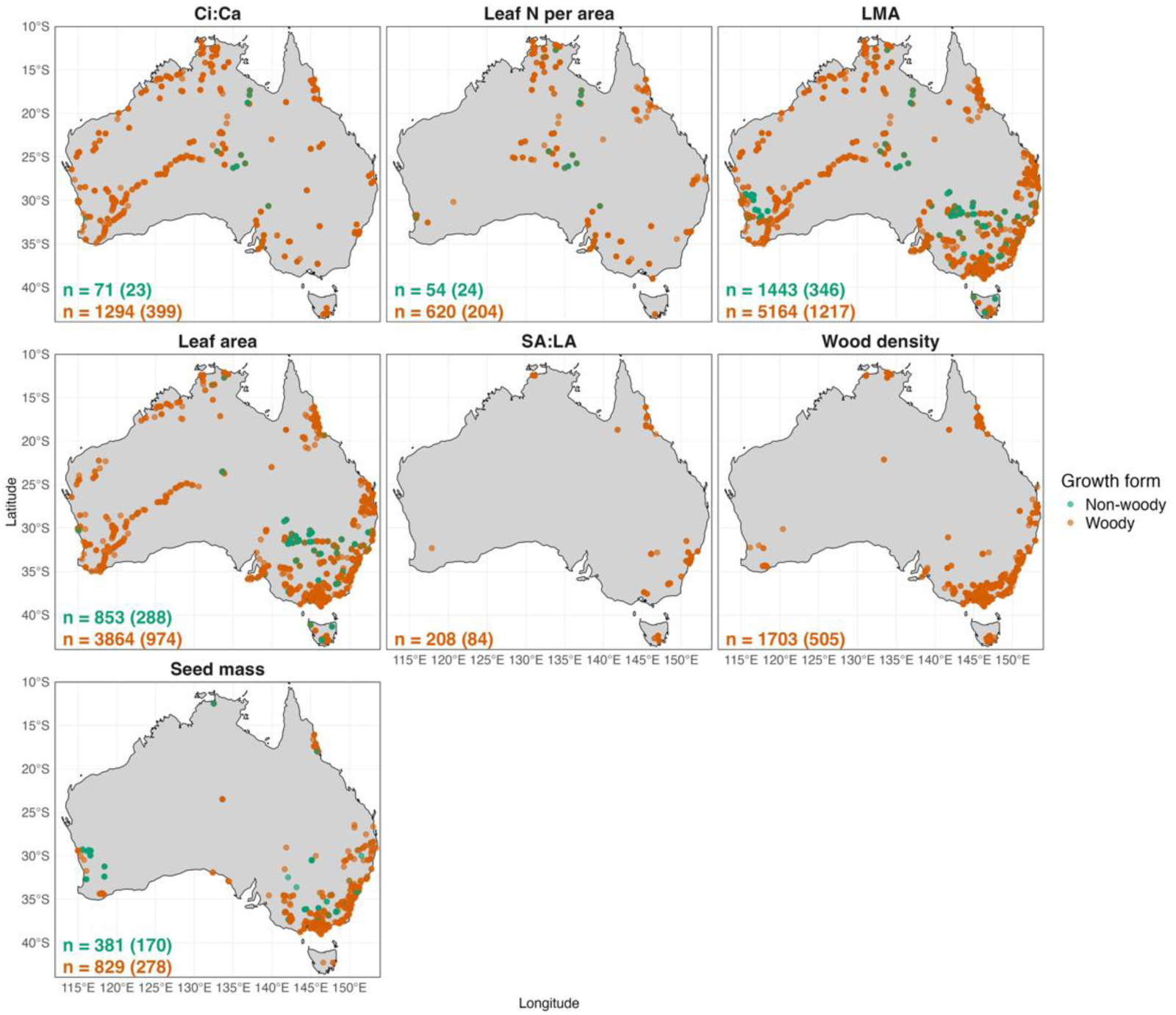
Distribution of trait data from AusTraits (Falster et al. 2021) used in the analysis. Brown points are from woody taxa while green points are from non-woody taxa. Data are filtered to taxa which have trait data recorded in two or more locations. The number of observations are presented for non-woody and woody taxa respectively, with the number of taxa in parentheses.

