## Supplementary material for "Species turnover dominates plant trait variation along continental climate gradients": Supp Tables S1-8 and Figures S1-5

### SUPPORTING INFORMATION

**Table S1.** The 0.1% and 99.9% quantiles of the species mean trait values per climate cell. Fold variation is the 99.9% quantile divided by the 0.1% quantile.

| Trait | Unit | 0.1% quantile | 99.9% quantile | Fold variation |
|---|---|---|---|---|
| Ci:Ca | unitless | 0.476 | 1.08 | 2.27 |
| Leaf N per area | g $m^{-2}$ | 0.56 | 21.3 | 38 |
| Leaf mass per area (LMA) | g $m^{-2}$ | 15.4 | 1,180 | 77 |
| Leaf area | $mm^2$ | 1 | 164,000 | 164,000 |
| SA:LA (Huber value) | $mm^2$ $cm^{-2}$ | 0.00551 | 0.507 | 92 |
| Wood density | g $cm^{-3}$ | 0.189 | 1.04 | 5.51 |
| Seed mass | mg | 0.00284 | 3,620 | 1,270,000 |

**Table S2**. Estimated $S_w$ and $\kappa_{obs}$ values and the correlation between $Env_m$ and $Env_w$ for each trait and environmental variable combination.

| | MAP | | | MAT | | |
|---|---|---|---|---|---|---|
| **Group** | **k (observed)** | **cor(Env_m , Env_w)** | **S_w** | **k (observed)** | **cor(Env_m , Env_w)** | **S_w** |
| **Ci:Ca** | | | | | | |
| Combined | 0.1006 | 0.02208 | 0.03282 [0.01317, 0.06183] | 0.1188 | 0.05081 | 0.08637 [0.0002287, 0.9230] |
| Woody | 0.1003 | 0.01782 | 0.02941 [0.01171, 0.05734] | 0.1221 | 0.04941 | 0.08596 [0.0002011, 0.9452] |
| Non-woody | 0.1275 | 0.1296 | 0.7384 [0.006919, 0.9993] | 0.05026 | 0.06876 | 0.5639 [0.001902, 0.9988] |
| **Leaf N per area** | | | | | | |
| Combined | 0.1328 | 0.05835 | 0.04814 [0.01409, 0.1066] | 0.07507 | 0.04491 | 0.02193 [0.00004489 , 0.6040] |
| Woody | 0.1295 | 0.06350 | 0.03705 [0.01099, 0.08099] | 0.07858 | 0.04242 | 0.02135 [0.00003645 , 0.5009] |
| Non-woody | 0.2399 | 0.05718 | 0.5533 [0.004026, 0.9983] | 0.04884 | 0.04334 | 0.5964 [0.003173, 0.9989] |
| **Leaf mass per area (LMA)** | | | | | | |
| Combined | 0.08457 | 0.003206 | 0.01751 [0.005424, 0.04088] | 0.1321 | -0.03006 | 0.7959 [0.2231, 0.9995] |
| Woody | 0.09606 | 0.002458 | 0.003637 [0.001453, 0.006783] | 0.1193 | -0.03390 | 0.2034 [0.03485, 0.9286] |
| Non-woody | 0.1345 | 0.008152 | 0.01600 [0.00005168, 0.1278] | 0.2139 | -0.01614 | 0.1231 [0.003599, 0.7966] |
| **Leaf area** | | | | | | |
| Combined | 0.08961 | 0.01071 | 0.00008747 [0.0000001969, 0.0008912] | 0.1525 | -0.02767 | 0.03416 [0.01823, 0.06058] |

**Table S2**. Estimated $S_w$ and $\kappa_{obs}$ values and the correlation between $Env_m$ and $Env_w$ for each trait and environmental variable combination.

| | MAP | | | MAT | | |
|---|---|---|---|---|---|---|
| **Group** | **k (observed)** | **cor(Env_m , Env_w)** | **S_w** | **k (observed)** | **cor(Env_m , Env_w)** | **S_w** |
| Woody | 0.09069 | 0.01082 | 0.0009649 [0.00006773, 0.003007] | 0.1621 | -0.03596 | 0.02484 [0.01283, 0.04423] |
| Non-woody | 0.1271 | -0.006361 | 0.02822 [0.0002282, 0.1443] | 0.1371 | 0.002120 | 0.6856 [0.05817, 0.9991] |
| **Seed mass** | | | | | | |
| Combined | 0.2197 | -0.08441 | 0.001034 [0.000002074, 0.01365] | 0.2066 | -0.06262 | 0.0008698 [0.000001837, 0.01084] |
| Woody | 0.2159 | -0.03617 | 0.003630 [0.000009326, 0.04226] | 0.2626 | -0.03801 | 0.0006989 [0.000001646, 0.007789] |
| Non-woody | 0.2525 | -0.1093 | 0.04789 [0.00009876, 0.9314] | 0.1736 | -0.08849 | 0.06364 [0.0001195, 0.9721] |
| **SA:LA (Huber value)** | | | | | | |
| Woody | 0.1612 | 0.01377 | 0.03189 [0.00007821, 0.3586] | 0.1805 | 0.003293 | 0.1394 [0.0003425, 0.9512] |
| **Wood density** | | | | | | |
| Woody | 0.1520 | -0.0001640 | 0.002256 [0.000009354, 0.01318] | 0.1135 | 0.04659 | 0.001910 [0.000003987, 0.02116] |

S_w: within-species share of climate-associated trait variation, median [2.5%, 97.5%] of the posterior at the observed k = var(Env_w) / var(Env_m).

**Table S3.** The overall responses of traits to either MAP or MAT based on ordinary least squares regression when all taxa are combined.

| | MAP | | | MAT | | |
|---|---|---|---|---|---|---|
| **Variable** | **Estimate** | **Std. error** | **p value** | **Estimate** | **Std. error** | **p value** |
| **Ci:Ca** | | | | | | |
| Intercept | 0.739 | 0.002 | <0.001 | 0.709 | 0.003 | <0.001 |
| Slope | 0.061 | 0.002 | <0.001 | 0.010 | 0.003 | <0.001 |
| **Leaf N per area** | | | | | | |
| Intercept | 0.860 | 0.019 | <0.001 | 1.052 | 0.028 | <0.001 |
| Slope | -0.328 | 0.019 | <0.001 | -0.113 | 0.021 | <0.001 |
| **Leaf mass per area (LMA)** | | | | | | |
| Intercept | 4.770 | 0.010 | <0.001 | 4.773 | 0.010 | <0.001 |
| Slope | -0.085 | 0.010 | <0.001 | -0.039 | 0.011 | <0.001 |
| **Leaf area** | | | | | | |
| Intercept | 6.454 | 0.027 | <0.001 | 6.546 | 0.030 | <0.001 |
| Slope | 0.994 | 0.026 | <0.001 | 0.555 | 0.032 | <0.001 |
| **SA:LA (Huber value)** | | | | | | |
| Intercept | -3.286 | 0.062 | <0.001 | -3.398 | 0.063 | <0.001 |
| Slope | -0.420 | 0.085 | <0.001 | -0.145 | 0.063 | 0.022 |
| **Wood density** | | | | | | |
| Intercept | -0.488 | 0.006 | <0.001 | -0.580 | 0.006 | <0.001 |
| Slope | -0.159 | 0.007 | <0.001 | -0.061 | 0.005 | <0.001 |
| **Seed mass** | | | | | | |
| Intercept | 0.444 | 0.057 | <0.001 | 0.777 | 0.064 | <0.001 |
| Slope | 0.601 | 0.077 | <0.001 | 0.742 | 0.065 | <0.001 |

**Table S4**. Parameter estimates and associated p values for the decomposition of the relationship between traits and mean annual precipitation (MAP; left column) and mean annual temperature (MAT; right column) into among- (orange; dashed lines) and within-species components (blue; solid lines).

| | MAP | | | MAT | | |
|---|---|---|---|---|---|---|
| **Variable** | **Estimate** | **Std. error** | **p value** | **Estimate** | **Std. error** | **p value** |
| **Ci:Ca** | | | | | | |
| Intercept | 0.742 | 0.003 | <0.001 | 0.719 | 0.006 | <0.001 |
| Across-species slope | 0.065 | 0.003 | <0.001 | 0.009 | 0.005 | 0.060 |
| Within-species slope | 0.038 | 0.007 | <0.001 | 0.008 | 0.007 | 0.311 |
| **Leaf N per area** | | | | | | |
| Intercept | 0.857 | 0.031 | <0.001 | 1.018 | 0.043 | <0.001 |
| Across-species slope | -0.326 | 0.032 | <0.001 | -0.082 | 0.037 | 0.026 |
| Within-species slope | -0.200 | 0.044 | <0.001 | -0.020 | 0.061 | 0.740 |
| **Leaf mass per area (LMA)** | | | | | | |
| Intercept | 4.784 | 0.018 | <0.001 | 4.784 | 0.019 | <0.001 |
| Across-species slope | -0.171 | 0.019 | <0.001 | -0.005 | 0.021 | 0.797 |
| Within-species slope | -0.078 | 0.017 | <0.001 | -0.077 | 0.015 | <0.001 |
| **Leaf area** | | | | | | |
| Intercept | 6.279 | 0.051 | <0.001 | 6.367 | 0.056 | <0.001 |
| Across-species slope | 1.060 | 0.052 | <0.001 | 0.660 | 0.061 | <0.001 |
| Within-species slope | 0.024 | 0.041 | 0.562 | 0.318 | 0.039 | <0.001 |
| **SA:LA (Huber value)** | | | | | | |
| Intercept | -3.335 | 0.080 | <0.001 | -3.442 | 0.083 | <0.001 |
| Across-species slope | -0.431 | 0.116 | <0.001 | -0.183 | 0.095 | 0.053 |
| Within-species slope | -0.179 | 0.220 | 0.412 | -0.169 | 0.146 | 0.237 |
| **Wood density** | | | | | | |
| Intercept | -0.478 | 0.010 | <0.001 | -0.571 | 0.009 | <0.001 |
| Across-species slope | -0.174 | 0.011 | <0.001 | -0.083 | 0.009 | <0.001 |
| Within-species slope | -0.021 | 0.015 | 0.156 | 0.009 | 0.013 | 0.513 |
| **Seed mass** | | | | | | |

| | MAP | | | MAT | | |
|---|---|---|---|---|---|---|
| **Variable** | **Estimate** | **Std. error** | **p value** | **Estimate** | **Std. error** | **p value** |
| Intercept | 0.541 | 0.095 | <0.001 | 0.809 | 0.109 | <0.001 |
| Across-species slope | 0.734 | 0.128 | <0.001 | 0.699 | 0.110 | <0.001 |
| Within-species slope | -0.008 | 0.074 | 0.918 | -0.001 | 0.065 | 0.989 |

SA:LA (Huber value) is from the Bayesian model: estimate = posterior mean, std. error = posterior SD, p value = two-sided posterior tail probability.

**Table S5**. Conditional effects of across-species (species turnover) and within-species variation in response to mean annual precipitation (MAP; left column) and mean annual temperature (MAT; right column) based on functional groups.

| | MAP | | | MAT | | |
|---|---|---|---|---|---|---|
| **Variable** | **Estimate** | **Std. error** | **p value** | **Estimate** | **Std. error** | **p value** |
| **Ci:Ca** | | | | | | |
| Across-species slope, non-woody | 0.009 | 0.013 | 0.517 | 0.009 | 0.013 | 0.471 |
| Across-species slope, woody | 0.067 | 0.003 | <0.001 | 0.009 | 0.005 | 0.081 |
| Within-species slope, non-woody | 0.053 | 0.042 | 0.212 | -0.033 | 0.068 | 0.630 |
| Within-species slope, woody | 0.037 | 0.007 | <0.001 | 0.007 | 0.007 | 0.317 |
| **Leaf N per area** | | | | | | |
| Across-species slope, non-woody | -0.152 | 0.141 | 0.281 | -0.010 | 0.128 | 0.936 |
| Across-species slope, woody | -0.350 | 0.032 | <0.001 | -0.097 | 0.038 | 0.011 |
| Within-species slope, non-woody | -0.386 | 0.251 | 0.124 | 0.519 | 0.391 | 0.184 |
| Within-species slope, woody | -0.191 | 0.043 | <0.001 | -0.039 | 0.061 | 0.529 |
| **Leaf mass per area (LMA)** | | | | | | |
| Across-species slope, non-woody | 0.197 | 0.048 | <0.001 | -0.116 | 0.048 | 0.017 |
| Across-species slope, woody | -0.454 | 0.015 | <0.001 | -0.043 | 0.020 | 0.029 |
| Within-species slope, non-woody | -0.070 | 0.049 | 0.152 | -0.095 | 0.040 | 0.018 |
| Within-species slope, woody | -0.088 | 0.016 | <0.001 | -0.062 | 0.016 | <0.001 |
| **Leaf area** | | | | | | |
| Across-species slope, non-woody | 0.504 | 0.100 | <0.001 | 0.068 | 0.091 | 0.453 |
| Across-species slope, woody | 1.058 | 0.062 | <0.001 | 0.758 | 0.074 | <0.001 |

| | MAP | | | MAT | | |
|---|---|---|---|---|---|---|
| **Variable** | **Estimate** | **Std. error** | **p value** | **Estimate** | **Std. error** | **p value** |
| Within-species slope, non-woody | -0.243 | 0.132 | 0.067 | 0.329 | 0.131 | 0.012 |
| Within-species slope, woody | 0.109 | 0.040 | 0.007 | 0.301 | 0.037 | <0.001 |
| **Seed mass** | | | | | | |
| Across-species slope, non-woody | 0.245 | 0.149 | 0.101 | 0.117 | 0.113 | 0.304 |
| Across-species slope, woody | 0.781 | 0.191 | <0.001 | 1.568 | 0.176 | <0.001 |
| Within-species slope, non-woody | 0.094 | 0.115 | 0.413 | -0.066 | 0.084 | 0.435 |
| Within-species slope, woody | -0.096 | 0.100 | 0.340 | 0.060 | 0.101 | 0.554 |

**Table S6.** Estimated AIC values for trait-environment models split by functional group with the estimated SD for the random intercept and slope either pooled or split across functional groups.

| | MAP | | | MAT | | |
|---|---|---|---|---|---|---|
| **Random-effect variances** | **AIC** | **ΔAIC** | **df** | **AIC** | **ΔAIC** | **df** |
| **Ci:Ca** | | | | | | |
| Both SDs pooled | **-3189.0** | **0.0** | **9** | -2802.6 | 10.5 | 9 |
| Slope SD by growth form | -3188.4 | 0.6 | 10 | -2802.5 | 10.7 | 10 |
| Intercept and slope SDs by growth form | -3188.7 | 0.3 | 11 | **-2813.1** | **0.0** | **11** |
| **Leaf N per area** | | | | | | |
| Both SDs pooled | 584.6 | 6.1 | 9 | 707.0 | 1.2 | 9 |
| Slope SD by growth form | **578.5** | **0.0** | **10** | **705.7** | **0.0** | **10** |
| Intercept and slope SDs by growth form | 580.2 | 1.7 | 11 | 707.7 | 2.0 | 11 |
| **Leaf mass per area (LMA)** | | | | | | |
| Both SDs pooled | 6949.7 | 197.6 | 9 | 7541.3 | 46.1 | 9 |
| Slope SD by growth form | 6915.8 | 163.7 | 10 | 7525.8 | 30.6 | 10 |
| Intercept and slope SDs by growth form | **6752.1** | **0.0** | **11** | **7495.1** | **0.0** | **11** |
| **Leaf area** | | | | | | |
| Both SDs pooled | 12575.6 | 56.0 | 9 | 12630.3 | 78.5 | 9 |
| Slope SD by growth form | 12525.1 | 5.6 | 10 | 12564.2 | 12.5 | 10 |
| Intercept and slope SDs by growth form | **12519.6** | **0.0** | **11** | **12551.7** | **0.0** | **11** |
| **Seed mass** | | | | | | |
| Both SDs pooled | 3407.1 | 2.8 | 9 | **3353.7** | **0.0** | **9** |
| Slope SD by growth form | 3408.5 | 4.1 | 10 | 3354.7 | 1.0 | 10 |
| Intercept and slope SDs by growth form | **3404.4** | **0.0** | **11** | 3355.9 | 2.2 | 11 |

AIC of the woody-split glmmTMB models (fixed effects identical) with the random-intercept and random within-species slope SDs pooled across growth forms, the slope SD estimated separately for woody and non-woody taxa, or both estimated separately. ΔAIC is relative to the best structure (bold) for each trait and climate variable; df = number of parameters.

**Table S7**. Estimated $S_w$ values when $\kappa$ is set to 1.

| | MAP | MAT |
|---|---|---|
| **Group** | **S_w** | **S_w** |
| **Ci:Ca** | | |
| Combined | 0.2522 [0.1171, 0.3958] | 0.4431 [0.001922, 0.9902] |
| Woody | 0.2320 [0.1056, 0.3775] | 0.4351 [0.001645, 0.9930] |
| Non-woody | 0.9568 [0.05182, 0.9999] | 0.9626 [0.03654, 0.9999] |
| **Leaf N per area** | | |
| Combined | 0.2757 [0.09713, 0.4732] | 0.2300 [0.0005977, 0.9531] |
| Woody | 0.2290 [0.07901, 0.4049] | 0.2173 [0.0004636, 0.9274] |
| Non-woody | 0.8377 [0.01657, 0.9996] | 0.9680 [0.06118, 0.9999] |
| **Leaf mass per area (LMA)** | | |
| Combined | 0.1741 [0.06058, 0.3351] | 0.9672 [0.6850, 0.9999] |
| Woody | 0.03661 [0.01492, 0.06637] | 0.6815 [0.2323, 0.9909] |
| Non-woody | 0.1079 [0.0003840, 0.5214] | 0.3963 [0.01661, 0.9482] |
| **Leaf area** | | |
| Combined | 0.0009753 [0.000002198, 0.009857] | 0.1883 [0.1086, 0.2973] |
| Woody | 0.01054 [0.0007463, 0.03219] | 0.1358 [0.07425, 0.2221] |
| Non-woody | 0.1860 [0.001793, 0.5703] | 0.9409 [0.3106, 0.9999] |
| **Seed mass** | | |
| Combined | 0.004690 [0.000009439, 0.05926] | 0.004196 [0.000008893, 0.05036] |
| Woody | 0.01660 [0.00004319, 0.1697] | 0.002657 [0.000006270, 0.02903] |
| Non-woody | 0.1662 [0.0003911, 0.9817] | 0.2814 [0.0006880, 0.9950] |
| **SA:LA (Huber value)** | | |
| Woody | 0.1696 [0.0004848, 0.7762] | 0.4730 [0.001894, 0.9908] |
| **Wood density** | | |
| Woody | 0.01466 [0.00006153, 0.08073] | 0.01659 [0.00003514, 0.1600] |

S_w: within-species share of climate-associated trait variation, median [2.5%, 97.5%] of the posterior at k = 1 for every group.

**Table S8.** The overall responses of traits to either MAP or MAT based on ordinary least squares regression when functional group is included as an interaction variable. Reported parameter estimates for Woody and Slope x woody are marginal estimates.

| | MAP | | | MAT | | |
|---|---|---|---|---|---|---|
| **Variable** | **Estimate** | **Std. error** | **p value** | **Estimate** | **Std. error** | **p value** |
| **Ci:Ca** | | | | | | |
| Intercept | 0.704 | 0.013 | <0.001 | 0.688 | 0.014 | <0.001 |
| Slope | 0.020 | 0.012 | 0.095 | 0.006 | 0.016 | 0.708 |
| Woody | 0.036 | 0.013 | 0.005 | 0.022 | 0.014 | 0.116 |
| Slope × woody | 0.042 | 0.012 | <0.001 | 0.004 | 0.016 | 0.802 |
| **Leaf N per area** | | | | | | |
| Intercept | 0.689 | 0.084 | <0.001 | 0.864 | 0.081 | <0.001 |
| Slope | -0.265 | 0.083 | 0.002 | -0.017 | 0.085 | 0.845 |
| Woody | 0.187 | 0.086 | 0.031 | 0.213 | 0.086 | 0.014 |
| Slope × woody | -0.074 | 0.086 | 0.390 | -0.109 | 0.088 | 0.215 |
| **Leaf mass per area (LMA)** | | | | | | |
| Intercept | 4.225 | 0.022 | <0.001 | 3.980 | 0.018 | <0.001 |
| Slope | 0.268 | 0.019 | <0.001 | -0.191 | 0.024 | <0.001 |
| Woody | 0.833 | 0.024 | <0.001 | 1.008 | 0.021 | <0.001 |
| Slope × woody | -0.673 | 0.021 | <0.001 | 0.143 | 0.026 | <0.001 |
| **Leaf area** | | | | | | |
| Intercept | 5.376 | 0.069 | <0.001 | 5.130 | 0.069 | <0.001 |
| Slope | 0.536 | 0.063 | <0.001 | 0.108 | 0.062 | 0.080 |
| Woody | 1.269 | 0.075 | <0.001 | 1.697 | 0.076 | <0.001 |
| Slope × woody | 0.421 | 0.069 | <0.001 | 0.480 | 0.071 | <0.001 |
| **Seed mass** | | | | | | |
| Intercept | -0.545 | 0.101 | <0.001 | -0.523 | 0.100 | <0.001 |
| Slope | 0.173 | 0.107 | 0.105 | 0.124 | 0.076 | 0.102 |
| Woody | 1.379 | 0.120 | <0.001 | 2.069 | 0.124 | <0.001 |
| Slope × woody | 0.493 | 0.150 | 0.001 | 1.364 | 0.119 | <0.001 |

**Figure S1.** Relationship between the mean average temperature and the mean temperature of the warmest quarter,the maximum temperature of the warmest month or the mean annual precipitation. In the top row, observations in blue are the cell-level climate values from the trait dataset used in analyses while observations in grey are the entire distribution of climate values in the Australian continent. The fitted lines are ordinary least squares linear regressions across either the data subset (blue) or the entire dataset (black). In the second row, the continental distribution of the mean average temperature or mean annual precipitation are plotted.

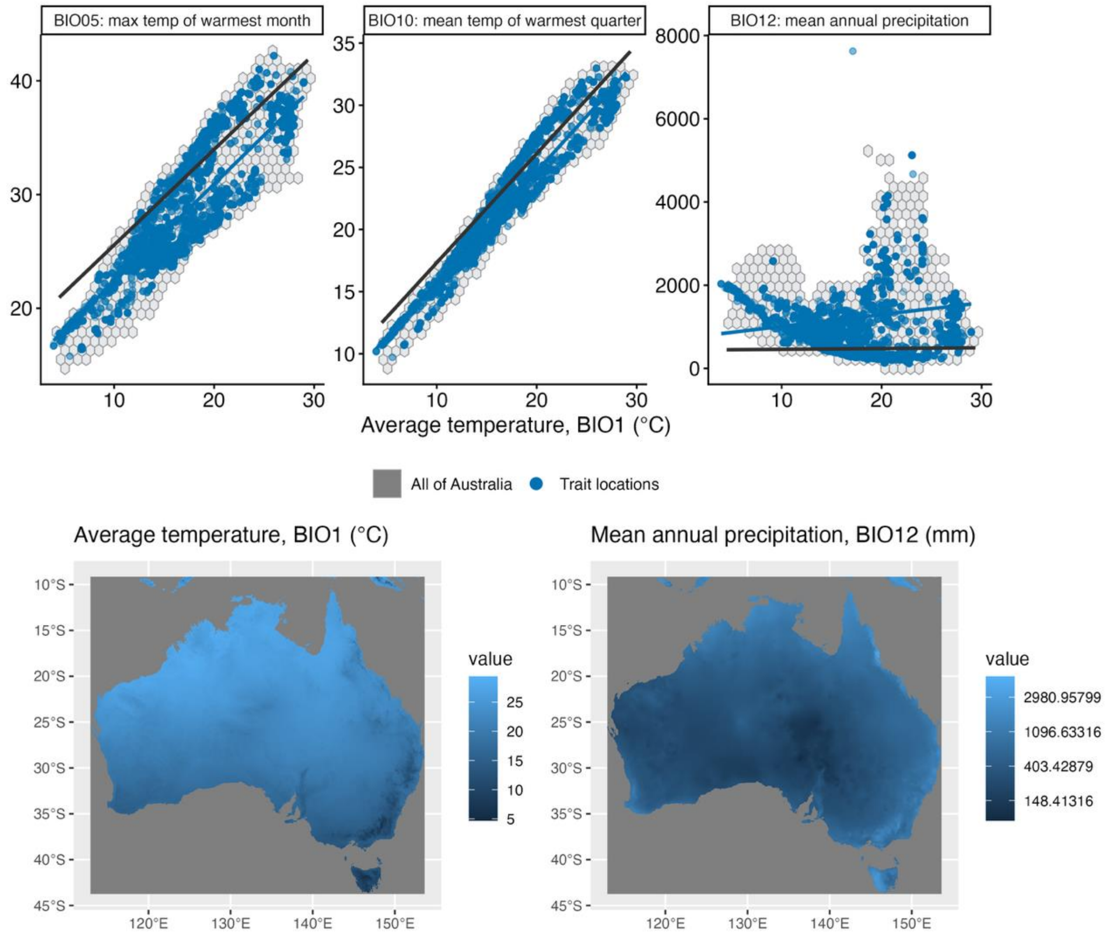

**Figure S2.** Density plots of the fitted species-level random slopes of traits with respect to deviation from species-level MAP centroids (i.e. the within-species response to MAP). The grey line is the fitted ordinary least squares regression of traits with respect to MAP. The orange line is the fitted slope for the response of traits to species-level MAP centroids. The dotted line is set at zero (i.e. no within-species response).

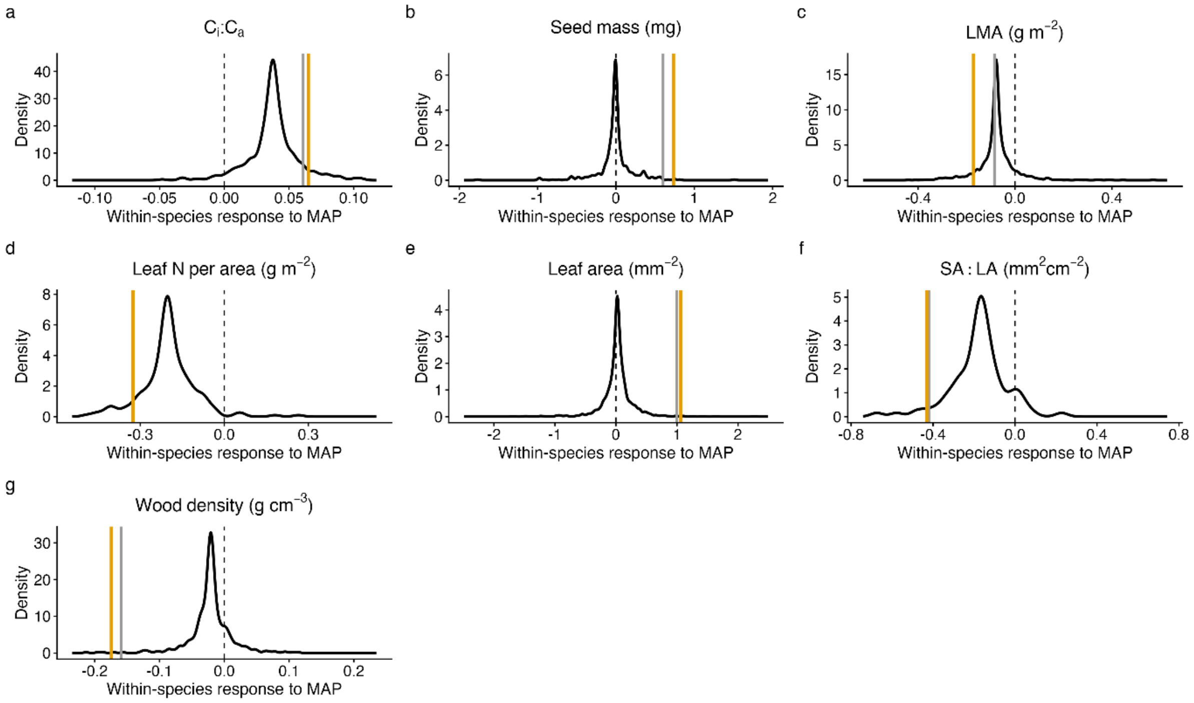

**Figure S3.** Density plots of the fitted species-level random slopes of traits with respect to deviation from species-level MAT centroids (i.e. the within-species response to MAT). The grey line is the fitted ordinary least squares regression of traits with respect to MAT. The orange line is the fitted slope for the response of traits to species-level MAT centroids. The dotted line is set at zero (i.e. no within-species response).

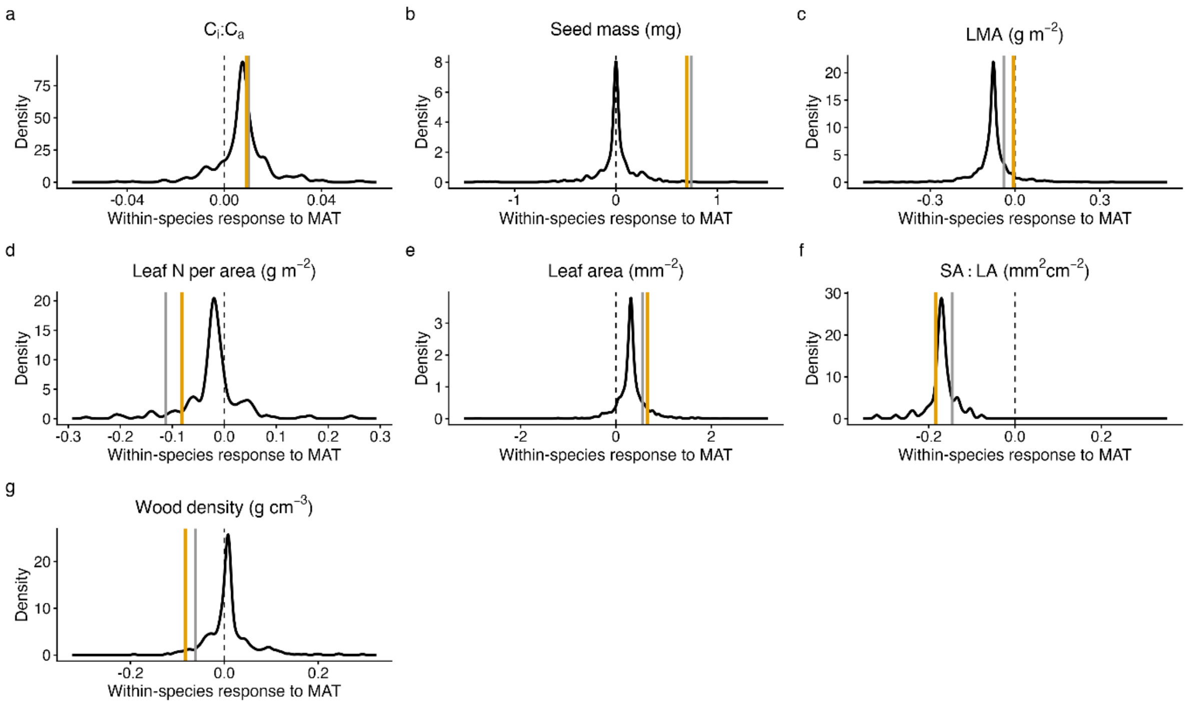

**Figure S4.** Within-species share of climate-associated trait variation $S_w$ along MAP and MAT gradients, evaluated at $\kappa = 1$ . $S_w$ is the fraction of the trait variation generated by the fitted model along each gradient that arises from within-species adjustment rather than species turnover, adjusted by a hypothetical $\kappa$ which is the climatic variation spanned within taxa relative to that spanned by taxon centroids, assumed here to be equal. Values of $S_w$ below 0.5 indicate that turnover generates more of the climate-associated variation than within-species adjustment. Points show posterior medians, thick bars 50% and thin lines 95% credible intervals, on a logit scale; intervals extending beyond the axis limits are truncated at the panel edge. Colours, white points and growth-form estimates as in Figure 2. Medians and 95% credible intervals are available in Table S2.

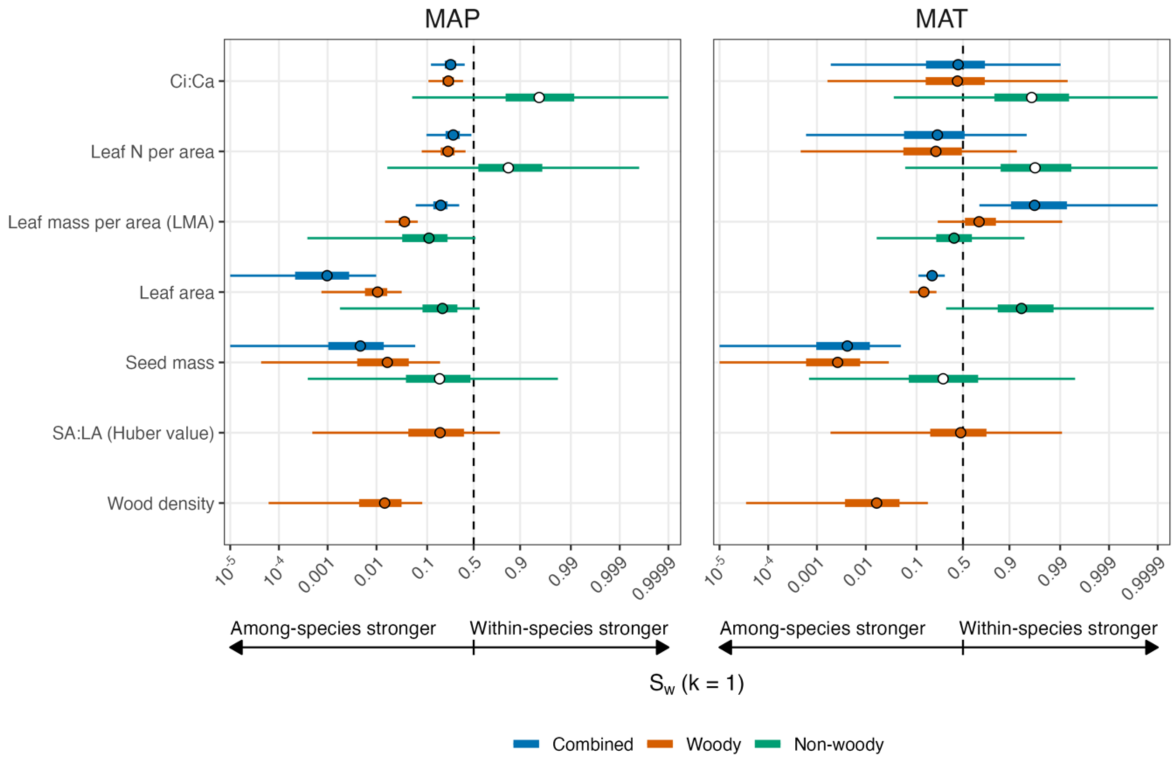

**Figure S5.** Sensitivity analysis of $S_w$ for each trait and environment combination with respect to simulated $\kappa$, ranging from $\kappa_{obs}$ to 1.

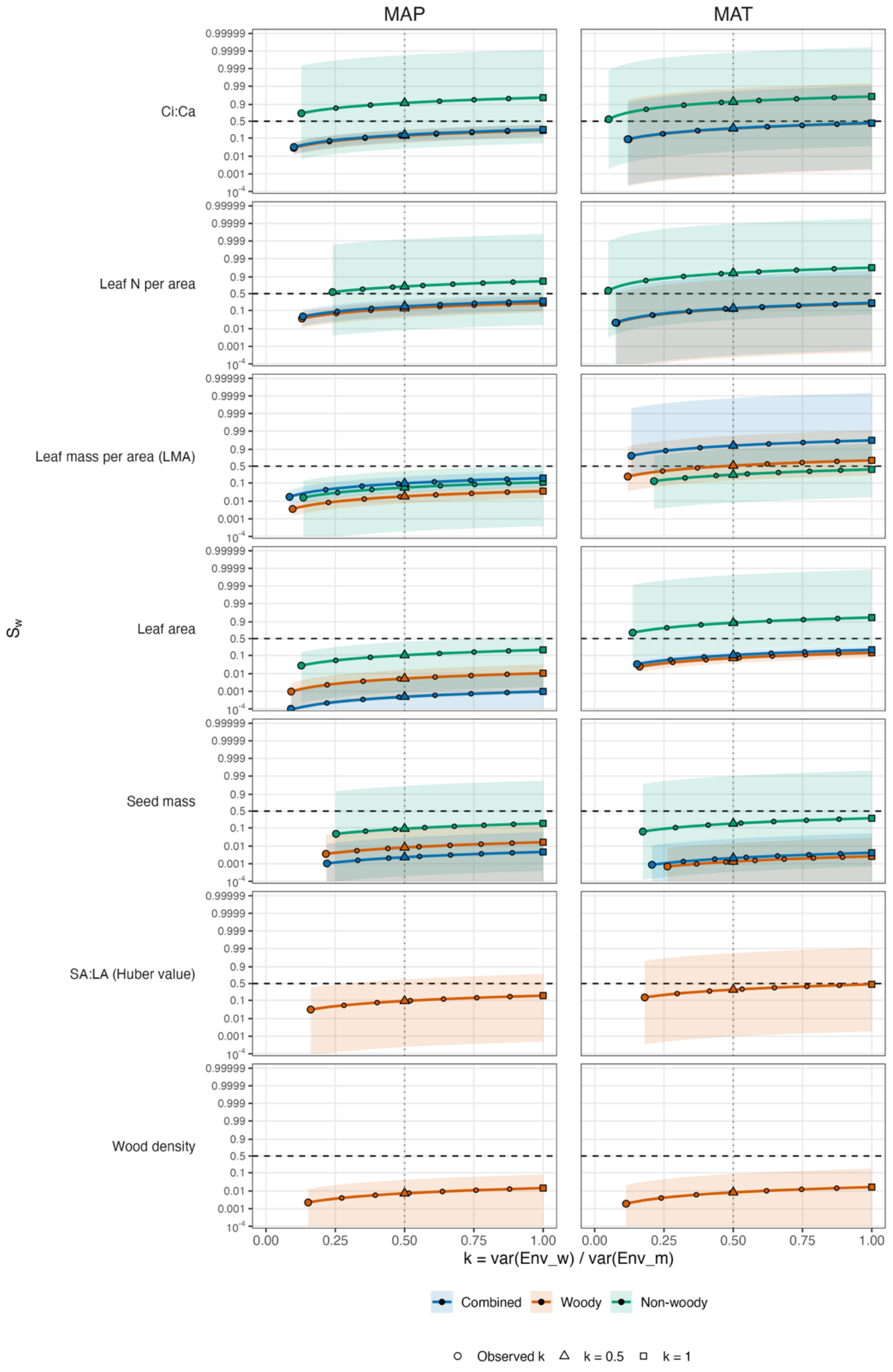
